# Structural basis for topological regulation of Tn3 resolvase

**DOI:** 10.1101/2021.12.07.471667

**Authors:** Sherwin P. Montano, Sally-J. Rowland, James R. Fuller, Mary E. Burke, Alasdair I. MacDonald, Martin R. Boocock, W. Marshall Stark, Phoebe A. Rice

**Affiliations:** University of Chicago, Chicago IL, USA; University of Glasgow, Glasgow, UK

## Abstract

Site-specific DNA recombinases play a variety of biological roles, often related to the dissemination of antibiotic resistance, and are also useful synthetic biology tools. The simplest sitespecific recombination systems will recombine any two cognate sites regardless of context. Other systems have evolved elaborate mechanisms, often sensing DNA topology, to ensure that only one of multiple possible recombination products is produced. The closely-related resolvases from the Tn3 and γδ transposons have historically served as paradigms for the regulation of recombinase activity by DNA topology. However, despite many proposals, models of the multi-subunit protein-DNA complex (termed the synaptosome) that enforces this regulation have been unsatisfying due to a lack of experimental constraints and incomplete concordance with experimental data. Here we present new structural and biochemical data that lead to a new, detailed model of the Tn3 synaptosome, and discuss how it harnesses DNA topology to regulate the enzymatic activity of the recombinase.

## Introduction

Tn3 resolvase belongs to a family of “small serine recombinases”. The serine recombinases are characterized by an active site serine residue which plays a key role in the strand exchange mechanism, attacking a backbone phosphodiester to break the DNA strand and becoming transiently covalently linked to one DNA strand end (reviewed in (Rice, 2015; Stark, 2014)(Johnson, 2015)). The small serine recombinases comprise an N-terminal catalytic domain which includes the active site, linked to a small C-terminal DNA-binding domain (herein referred to as DBD). A separate group of ‘large serine recombinases’ which generally serve as transposases or bacteriophage integrases have much larger C-terminal extensions and utilize different regulatory mechanisms (Smith, 2015).

Tn3 resolvase is encoded by the penicillin-resistance-carrying Tn3 transposon (Arthur and Sherratt, 1979; Heffron et al., 1979). Replicative transposition of Tn3 leads to a “cointegrate” product in which the donor and recipient replicons are fused, with copies of the transposon at each junction oriented in direct repeat (Figure 1a). Resolvase then efficiently separates the replicons by promoting recombination between the two copies of Tn3 at a specific 114-bp “*res*” site.

**Figure 1.**
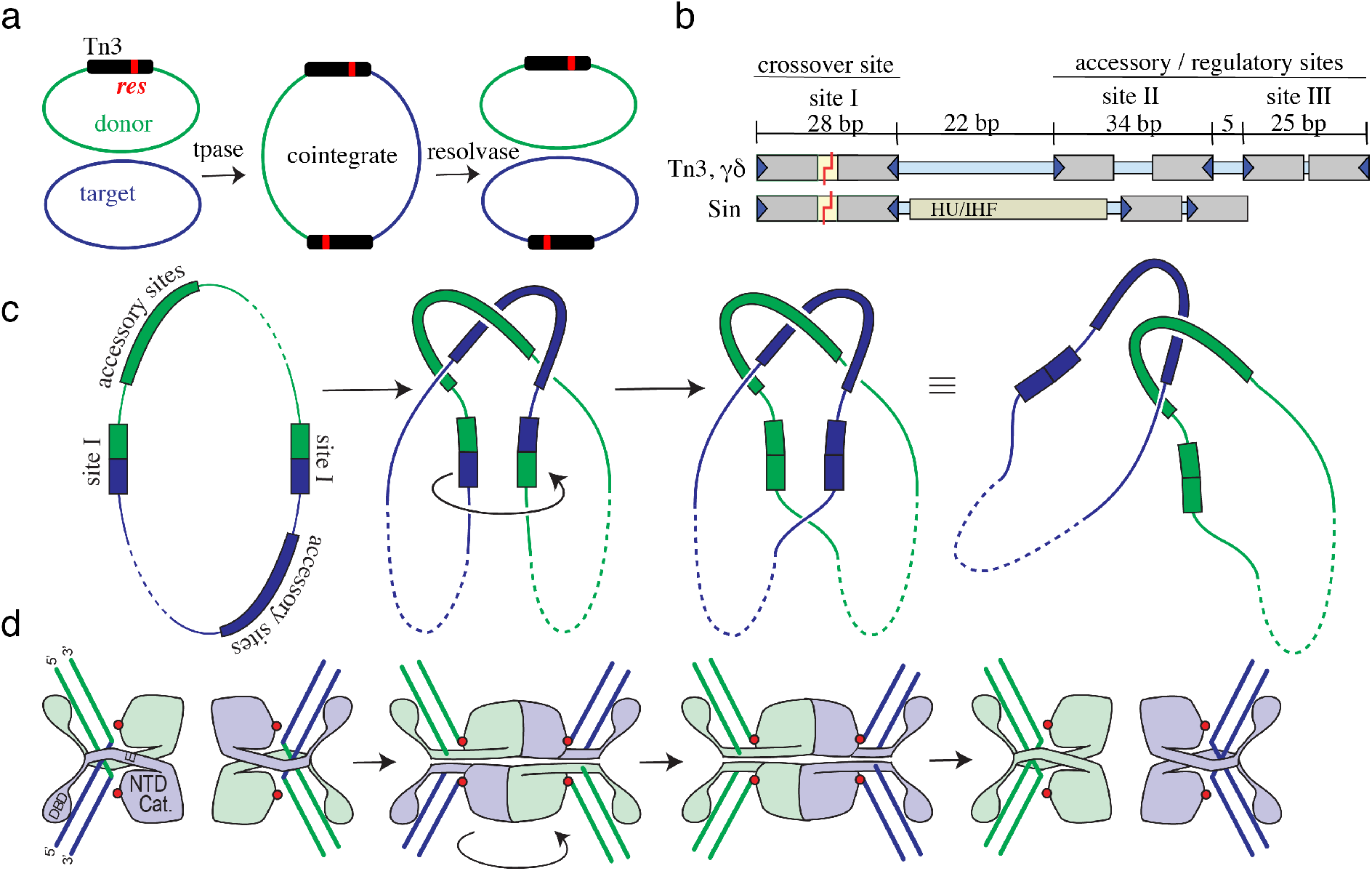
Function of Resolvase. a) Replicative transposition of the Tn3 transposon (black bar) creates a cointegrate product that is resolved by Tn3 resolvase acting at the *res* sites (red) within each copy of the transposon. b) Tn3/γδ and Sin *res* sites. Binding sites for individual resolvase subunits are shown as gray bars with triangles indicating motifs at the ends of the binding sites recognized by the resolvase C-terminal DNA-binding domains (DBDs). The *res* sites for the Tn3 and γδ transposon resolvases differ slightly in sequence, but the spacing and orientations of the 3 dimer-binding sites are identical and the proteins are functionally interchangeable (Grindley et al., 1982). The *res* site for Sin contains only 2 dimer-binding sites but includes a site for a DNA bending protein (HU or IHF)(Rowland et al., 2006). c) DNA transactions during recombination. Formation of the synaptic complex by interactions of the proteins bound to the two *res* sites traps 3 supercoiling nodes and juxtaposes the two crossover sites (*res* binding site I). Supercoil nodes in the 1^st^ panel and those outside of *res* are not drawn in the diagrams, for simplicity. Within this complex, cleavage of both DNA strands in both crossover sites is followed by a 180° rotation of one pair of DNA half-sites relative to the other pair. Religation of the ends completes recombination, and the cointegrate is converted into two catenated DNA circles (which can be separated from each other by a type II topoisomerase from the host). d) Cartoon showing resolvase actions at site I. Resolvase binds isolated site Is as dimers, but when incorporated into the synaptosome (2^nd^ panel) undergoes a large activating conformational change to form a tetramer with a flat central interface. Each subunit’s active site serine residue (S10; red dots) then attacks the DNA, creating double-strand breaks with 2-nt 3’ overhangs, and with a phosphoserine linkage covalently attaching each broken end to a subunit. A 180° rotation of the lower two subunits (relative to the rest of the synaptosome) realigns the broken ends, which are resealed by the attack of the 3’OH groups on the phosphoserine linkages.

The Tn3 *res* site carries three different dimer-binding sites, all of which are required for efficient resolution (Figure 1b and Figure 1supplement 1). Only the subunits bound at site I can be catalytically active, and only when incorporated into the “synaptosome”, the complex of two *res* sites bound by six resolvase dimers. The dimers bound to binding sites II and III play non-catalytic, “architectural” roles, assembling the scaffolding of the synaptosome. Once juxtaposed within the synaptosome, the two site I-bound resolvase dimers undergo a large conformational change leading to a catalytically active tetramer (Figure 1c,d). The active site serine of each subunit within the tetramer then attacks the phosphodiester backbone, becoming covalently linked to the 5’ end and displacing a DNA 3’OH, thus generating double-strand breaks with 2-nt 3’ overhangs. A 180° rotation of two subunits relative to the other two then realigns the DNA ends, and the cleavage reaction is reversed to form the ligated recombinant product (Figure 1c,d) (reviewed in (Rice, 2015; Stark, 2014)).

The synaptosome acts as a topological filter to ensure that recombination can only proceed between appropriately paired sites. Detailed studies of the DNA topological changes accompanying recombination of a standard substrate (a supercoiled plasmid with directly repeated *res* sites) by Tn3 and γδ resolvases determined that the two *res* sites wrap around one another, trapping 3 negative supercoiling nodes within the synaptosome (Figure 1c). This wrapping requirement prevents incorrect alignment of the two *res* sites, which could result in inversion of the DNA segment between the sites rather than resolution, or synapsis of *res* sites on separate replicons, which could lead to unwanted replicon fusions. The topology of the synaptosome also allows energy stored as torsional strain (negative supercoiling) in the DNA to drive the reaction forward (Grindley et al., 2006; Stark, 2014).

Structural studies of several different small serine recombinases have elucidated the dimer and tetramer conformations (cartooned in Figure 1d). The N-terminal catalytic domain is connected by a long helix (helix E) and a flexible segment to the C-terminal helix-turn-helix DNA-binding domain (DBD). The WT proteins always crystallize in the inactive dimeric conformation (Mouw et al., 2008; Rice and Steitz, 1994a; Sanderson et al., 1990; Yang and Steitz, 1995), but numerous constitutively active resolvase mutants have been isolated that can form tetramers in the absence of the architectural portion of the synaptosome (e.g. (Sarkis et al., 2001) (Mouw et al., 2010; Olorunniji et al., 2008)(Sanders and Johnson, 2004)(Dhar et al., 2009)). Tetramerization entails both intra- and inter-molecular repacking of helix E. Slicing through the tetramer is a large, flat hydrophobic interface about which rotation occurs. Supporting the rotational mechanism for strand exchange, structures of tetramers in multiple rotational states have been determined (Kamtekar et al., 2006; Keenholtz et al., 2011; Li et al., 2005; Trejo et al., 2018)(Ritacco et al., 2013).

Until now only one small serine recombinase, the staphylococcal Sin resolvase, has been crystallized bound to accessory site (‘architectural’) DNA (Mouw et al., 2008). Sin and Tn3 resolvases are quite distantly related, and their *res* sites also differ significantly (Figure 1b). The Sin *res* has only one accessory site (site II) that binds a Sin dimer. The Sin – site II structure is highly asymmetric and would have been very difficult to predict based solely on existing structures of γδ resolvase bound to site I DNAs. The inter-dimer contacts seen in the crystal were shown by independent genetic screens and biochemical experiments to be critical for synaptosome formation (Rowland et al., 2009). This structure provided the missing puzzle piece that, together with structures of γδ resolvase and IHF bound to DNA, allowed modeling of the full Sin synaptosome (Mouw et al., 2008). However, although one similar inter-dimer contact was known to be important for Tn3/γδ synaptosome formation, the Sin model could not be extrapolated to the Tn3 synaptosome because the architectural portions of the two *res* sites are very different (Figure 1b).

As noted above, elucidation of the structure of the Tn3/γδ synaptosome has been a longstanding challenge, and two major issues have blocked progress. The first issue has been a lack of structural information for Tn3/γδ resolvase bound to sites other than site I. The second issue has been incomplete understanding of the dimer-dimer interactions that hold the synaptosome together. It has long been known that a small group of catalytic domain residues mediate an important dimer-dimer contact, historically termed the “2-3’ interaction” and now simply the *R* (for regulatory) interface (Hughes et al., 1990; Rowland et al., 2009). The *R* interface was observed in the first structures of γδ resolvase in the absence of DNA, and later in the Sin – site II complex structure (Mouw et al., 2008; Sanderson et al., 1990). Mutations of *R* interface residues abolish recombination activity, presumably by disrupting critical interactions within the synaptosome (Hughes et al., 1990; Murley and Grindley, 1998). However, because all 12 protein subunits in the synaptosome are identical in sequence, determination of the *R* interface contacts of specific subunits has been very difficult. Recently we used hybrids of Tn3 resolvase and a related resolvase with different DNA-binding and *R*-interface specificities to identify individual dimer-dimer interactions, and to show that the only essential interactions of the site II- and III-bound dimers are via the *R* interface (Rowland et al., 2020). However, that work still did not determine the specific network of pairwise connections between individual subunits.

Here we report a series of structural and biochemical studies leading to a 3D model of the Tn3 resolvase synaptosome. We determined the structures of the resolvase-site III complex in two different space groups and then used those structures to create a model of the resolvase – site II complex that is compatible with SAXS and DNA footprinting data. We have also developed our toolbox of hybrid resolvases and synthetic *res* sites to pinpoint interactions between specific subunits within the synaptosome. Our new model of the synaptosome is consistent with these data, as well as previously published biochemical studies.

The new model suggests that activation of the crossover site – bound subunits may involve a mechanical form of allosteric activation mediated by the *R* interface, as well as a high local concentration of proteins (see Discussion). The Tn3 resolvase structures presented here, together with our previous Sin structures, also illuminate how the same protein can serve both catalytic and architectural roles, with the length and geometry of the binding sites defining the conformation of the DNA-bound dimers, and hence the topology of recombination.

## Results

### Tn3 resolvase dimer – site III structure

Tn3 resolvase - site III complexes were crystallized in two different space groups. Form I contained 1 complex per asymmetric unit, diffracted with minor anisotropy to ~3.4Å, and was phased by MIRAS (multiple isomorphous replacement with anomalous scattering). Form II contained two complexes per asymmetric unit and diffracted more anisotropically to ~ 4 /4.5 /4Å along the 3 principal axis, respectively. A partial model was obtained by molecular replacement, after which the map was improved by combining SIR (single isomorphous replacement)- and model-derived phases. While the positions of the DNA and protein secondary structure elements were clear, density for many of the side chains was poor. Modeling of the amino acid sequence register and side chain placement was facilitated by comparison with structures of γδ resolvase (Rice and Steitz, 1994a; Yang and Steitz, 1995). Refinement and data processing statistics are listed in Table 1. In both crystal forms the DNA duplexes pack end-to-end forming pseudo-continuous helices that extend throughout the crystal and are stabilized by base pairing between 1-nt overhangs in form 1 and 2-nt overhangs in form II. The 3 independent determinations of the Tn3 resolvase dimer – site III structure provided by these two crystal forms overlay quite well (Figure 2), with minor differences in the DNA bending angle.

**Figure 2.**
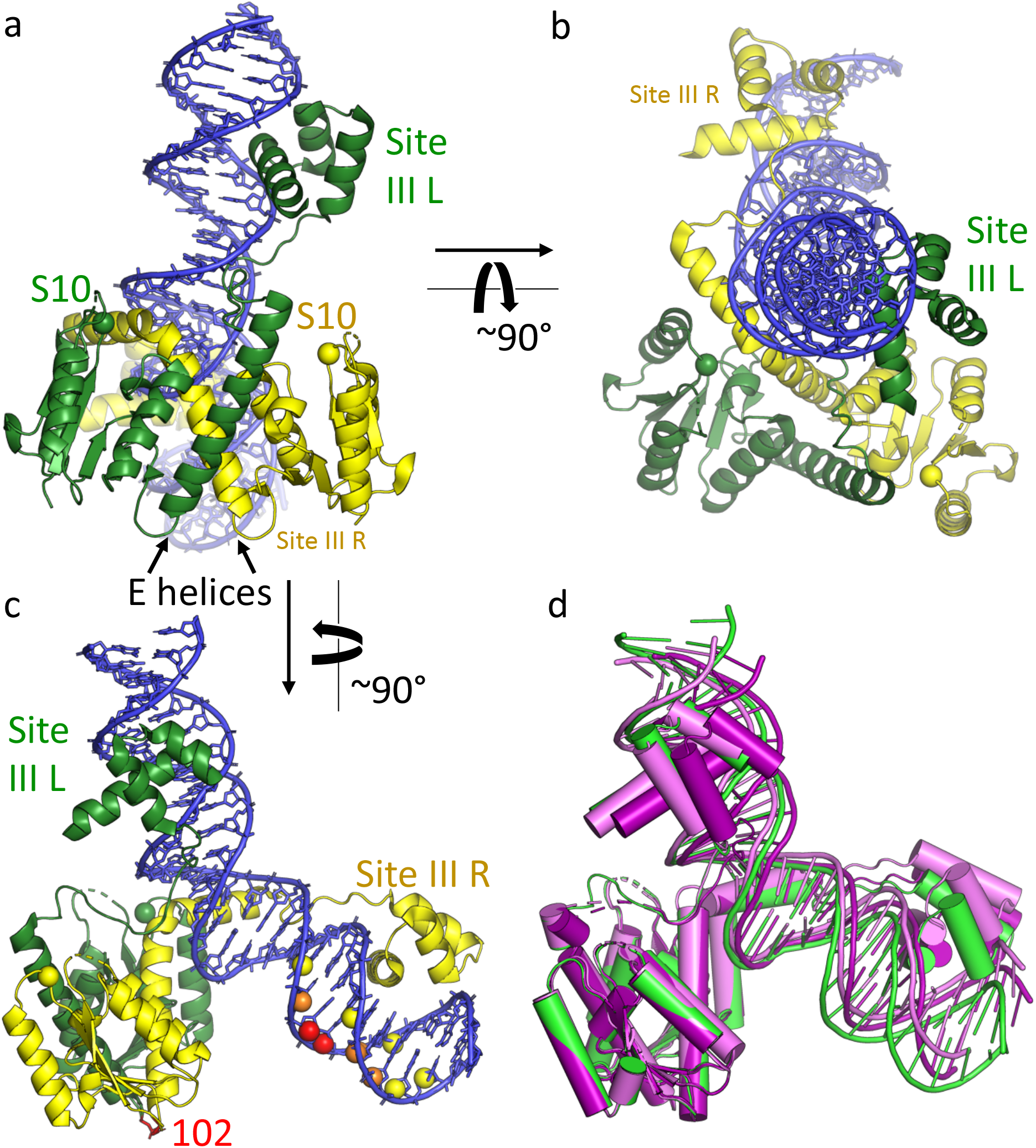
Structure of the Tn3 resolvase - site III complex. a,b,c) orthogonal views. Green: left half-site bound subunit, yellow, right half-site bound subunit. The alpha-carbons of the active site serines are shown as spheres. In (c) the targeted cleavage data of (Mazzarelli et al., 1993) are mapped onto the structure. Residue 102 of both subunits, where EDTA-iron complexes were tethered, is shown as red sticks, and the C4’ atoms in the DNA are colored according to the intensity of the resulting cleavage at each nucleotide. d) Superposition of the 3 independent determinations of the complex provided by the 2 crystal forms, using the N-terminal catalytic domain dimers as guides. Green, crystal Form I; pale and bright pink, Form II.

The complex is highly asymmetric, with the pseudo-twofold axis relating the two DNA binding domains roughly perpendicular to that between the two catalytic domains (Figure 2). Helix E of the subunit that binds the right half-site (site IIIR, as drawn in Figure 1b) is kinked by ~80°, with its C-terminal segment inserted into the minor groove at the center of the site. This leaves no space for the C-terminal segment of the other subunit’s helix E, which is very poorly ordered and may be partially unfolded. The asymmetric positioning of the catalytic domains in our structures agrees well with previous targeted DNA cleavage experiments (Figure 2c)(Mazzarelli et al., 1993). The placement of the catalytic domain dimer appears to be stabilized by interactions with the side of helix E that faces away from the DNA. Three negatively charged residues (E124, E128 and E132) extend from one subunit’s E helix toward the positively charged active site region of the other subunit’s catalytic domain, where DNA interacts in the site-I bound catalytic tetramer (e.g. 1ZR4; (Li et al., 2005)). The central of these residues, E128, was identified in a recombination-defective mutant (E128K) with a specific defect in site III binding. (Grindley, 1993; Hatfull et al., 1987; Rowland et al., 2020).

That the right rather than the left subunit’s E helix is consistently the one docked into the minor groove, despite the overall symmetry between DBD-binding motifs within site III, shows that the “spacer” sequence in the center of site III is by no means random, and that the C-terminal segment of helix E does have DNA sequence preferences. As in the γδ resolvase – site I complex structure, most of the DNA bending occurs at a central kink where there is a large roll angle between adjacent base pairs, and T126 from one of the E helices is partially intercalated. The roll angles in the Tn3 complexes range from 33° to 38°, and the overall DNA bend angles from ~42° – 50°. Once the stacking at the central kink has been disrupted, the energy barrier to further bending is probably low, allowing flexibility in the overall bend angle.

Comparison to other serine recombinase dimer– DNA complex structures shows that the arrangement of the DNA half-sites and the spacer sequence dictates the geometry of the overall complex in those cases as well. In all three structures shown in Figure 3 the docking of helix E into the minor groove widens it, bending the DNA away from the protein. While at least one of the E helices is kinked in every case, the degree and direction of the kink vary widely, and the bound DNAs are almost mutually perpendicular. Sin’s site II also forces a mismatch in the symmetry between the DBDs and the catalytic domains, in this case because the half-sites in the DNA are in direct rather than inverted repeat. As with Tn3 site III, only one of the E-helices can be accommodated in the minor groove near the center of site II. In contrast, the γδ resolvase - site I complex is only slightly asymmetric. The site I central spacer is 3 bp longer than that of site III, providing room for both E-helices to dock into the minor groove, although one is still slightly kinked.

**Figure 3.**
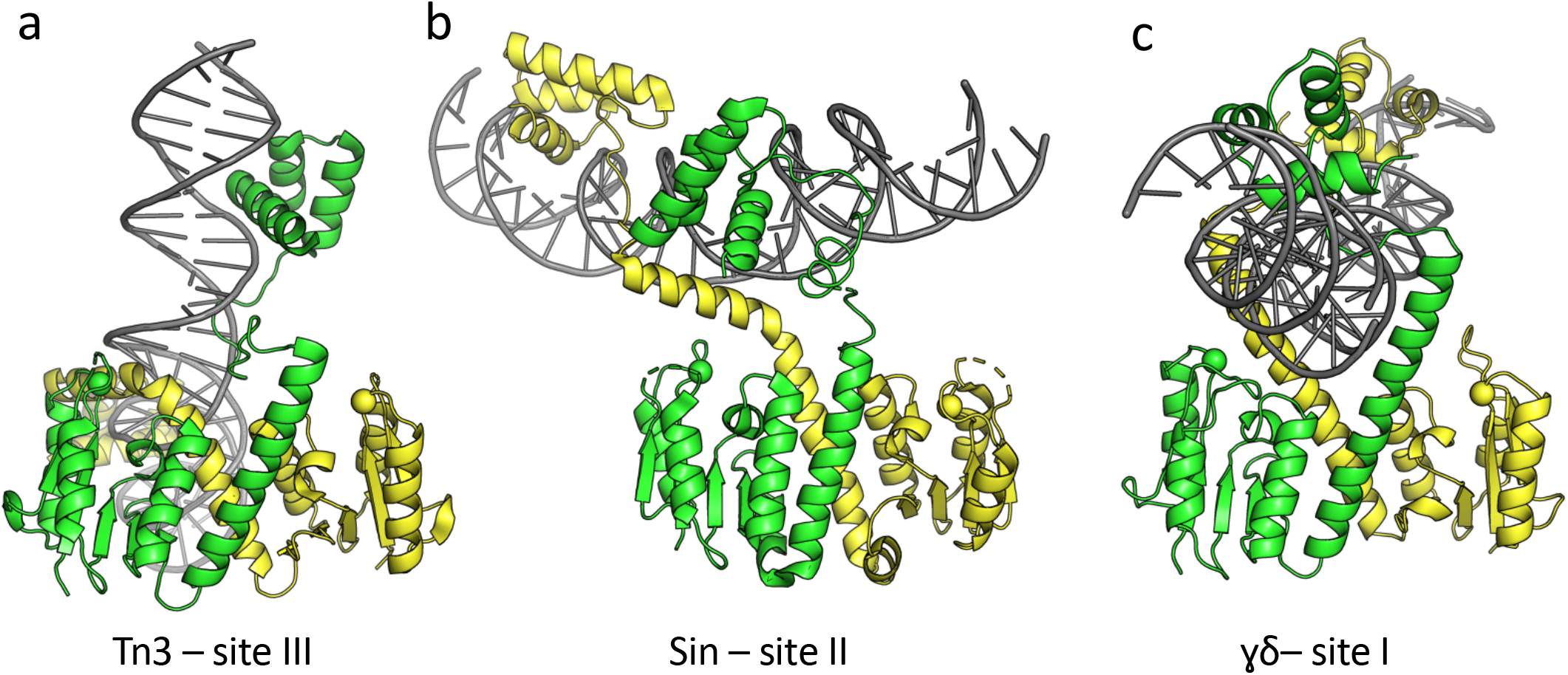
Comparison of serine resolvase – DNA complex structures. In each panel, the catalytic domain dimers are oriented similarly, and the Cα’s of the active site serines are highlighted as spheres. a) Tn3 resolvase bound to its cognate site III. b) Sin resolvase bound to its cognate site II. (PDB id 2r0q; (Mouw et al., 2008)) c) γδ resolvase bound to the crossover site, site I. (PDB id 1gdt; (Yang and Steitz, 1995))

### Conserved inter-dimer contacts

In Form II crystals, the catalytic domains of the two dimers in the asymmetric unit interact via a patch of charged residues on the corner of the protein that is farthest from the DNA and the dimer interface, very similarly to the previously observed *R* interface (see Introduction and Figure 4).

**Figure 4.**
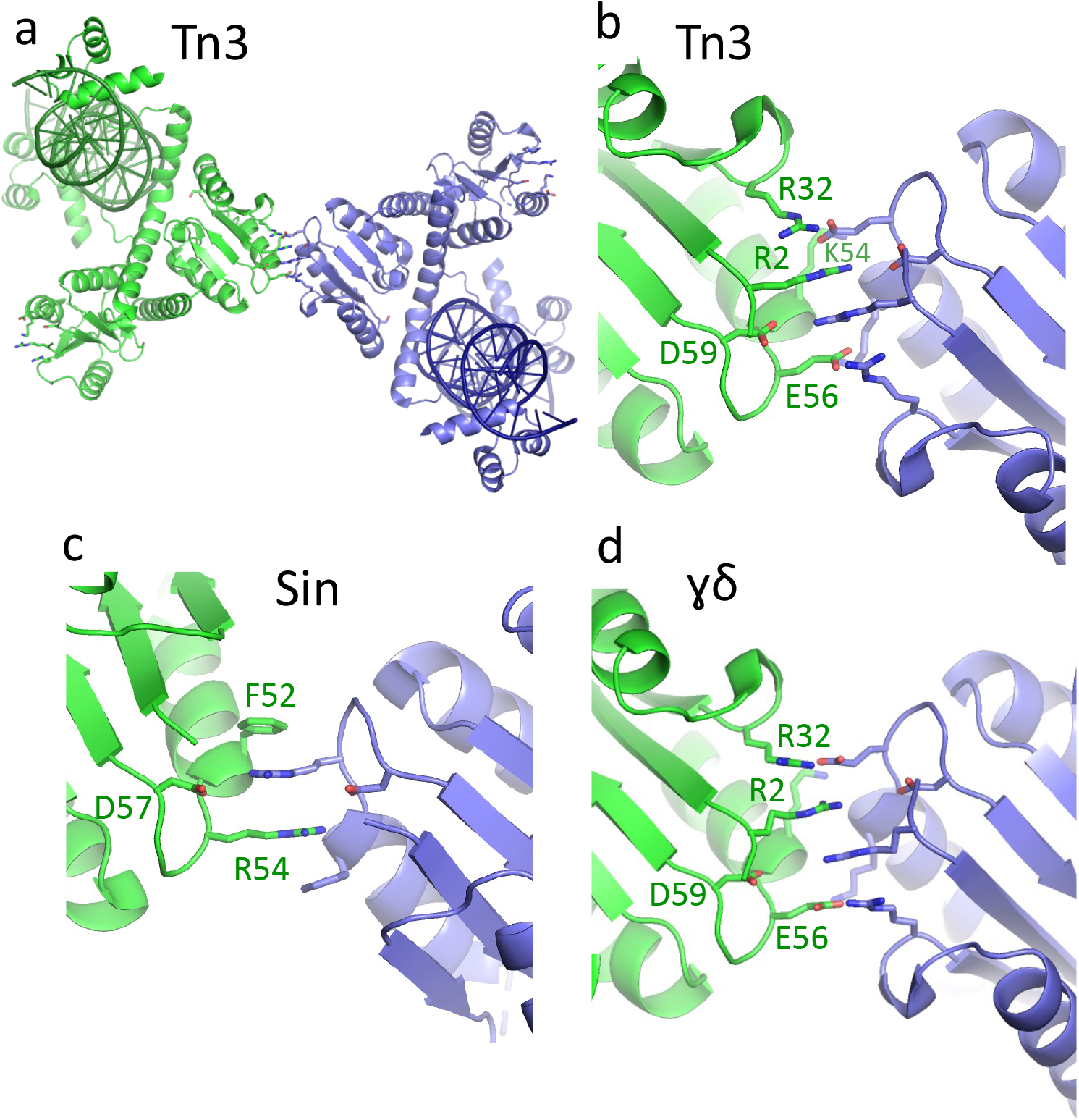
The *R* interface is conserved. a) Packing of the two Tn3 resolvase – site III bound dimers in crystal form II, with side chains known to be important for the *R* interface shown as sticks. b) Closeup of the interactions at the center of panel (a). Due to the low resolution of this crystal form, side chain details are uncertain. c) The *R* interface seen in the crystal between Sin – site II bound dimers. (PDB id 2r0q; (Mouw et al., 2008)) d) The *R* interface seen in the first serine recombinase structure, that of the γδ resolvase catalytic domain. (PDB id 2rsl; (Rice and Steitz, 1994a))

The Sin – site II structure revealed surprisingly extensive contacts between the DBDs of two dimers, and DBD-mediated interactions between site II-bound Sin dimers are required for synaptosome formation (in addition to *R*-interface-type contacts between dimers bound at site I and site II). We therefore examined the crystal packing interactions in our Tn3 – site III structures, and found a DBD-mediated dimer-of-dimers in crystal Form I. However, the contact surfaces involved are much smaller than for Sin and these interactions are not recapitulated in Form II. Furthermore, our recent functional experiments show that DBD-DBD contacts are not required for Tn3 resolvase synaptosome formation (Rowland et al., 2020).

### Modeling site II

The spacer between DBD-binding motifs in site II is 9 bp (nearly one helical turn) longer than that in site III (see Figure 1 supplement 1b). However, the DNA sequence of site IIL is quite similar to that of IIIR, and targeted DNA cleavage experiments implied that site II also forms an asymmetric complex with resolvase, but with the left rather than the right half-site more closely associated with the catalytic domains (Mazzarelli et al., 1993). We therefore modeled the site II-bound complex based on our site III-bound crystal structures by flipping the orientation, then adding 9 bp of DNA to the inner portion of site IIR, using the crystal structure of an A-tract containing duplex to model the run of A-T basepairs in this segment that is conserved in Tn3-like *res* sites (see Materials and Methods for more detail). Fully unfolding and extending the C-terminal segment of helix E allows the site IIR-bound DBD to dock as in all other half-sites (Figure 5a). The center of the bend predicted by this model matches that predicted earlier from circular permutation binding experiments (Blake et al., 1995). To fit this model into the full synaptosome model described below, the magnitude (but not the direction) of the central bend was increased, and a small additional bend was introduced at the A-tract – site IIR junction.

**Figure 5.**
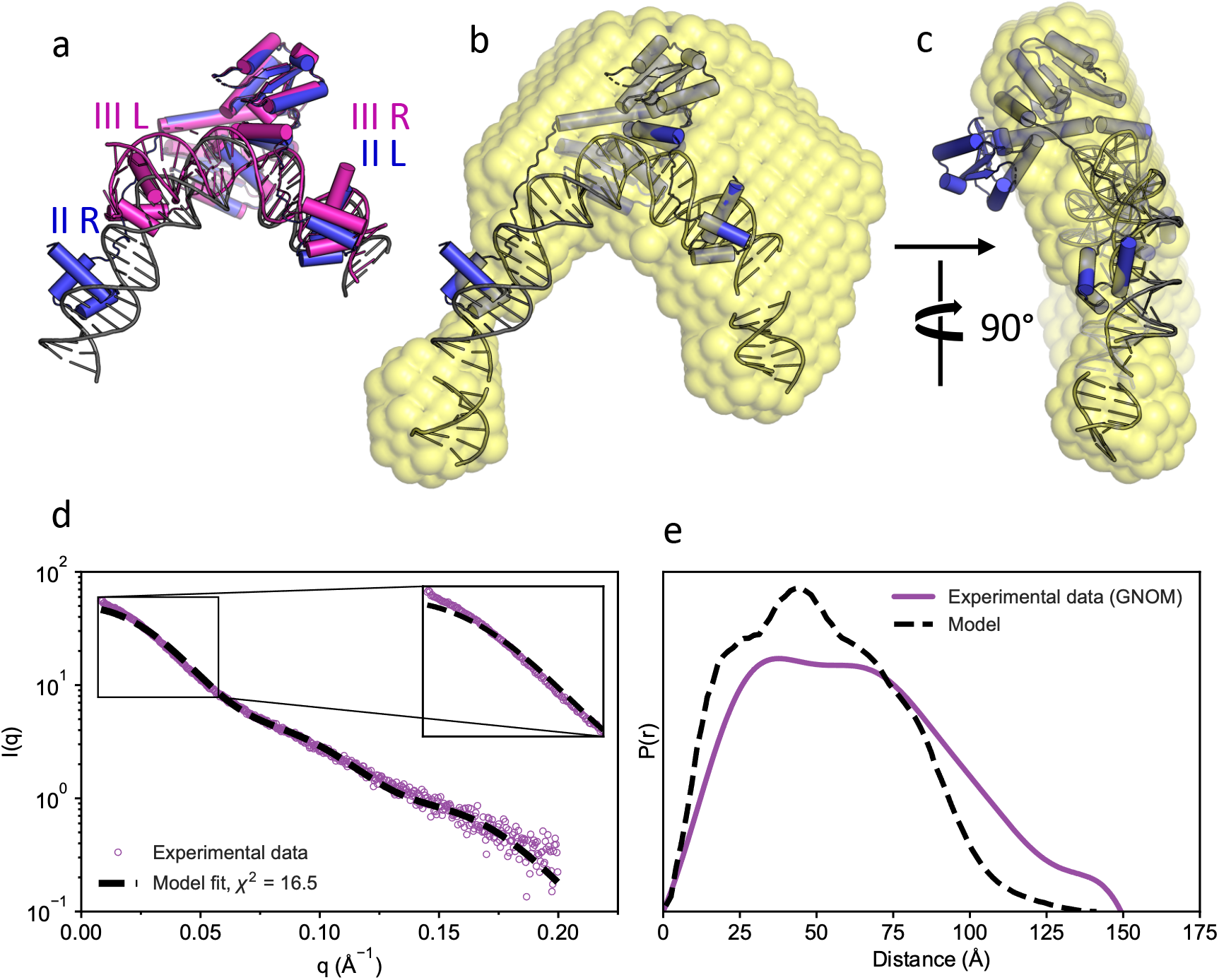
Modeling the Tn3 resolvase - site II complex. a) The site II – bound model (blue and gray) is superimposed onto the site III-bound structure (pink). The catalytic domains were aligned such that site IIIR and site IIL are superimposed. b) A surface representation of the SAXS bead model is shown in transparent yellow, manually superimposed onto the site II-bound model (blue and gray). Adjacent DNA segments from the synaptosome model are also shown so that the DNA duplex displayed is exactly the same length as that used in the SAXS experiment. c) Orthogonal view of part (b). d) The calculated SAXS profile from the atomic model shown in **(b)** and **(c)** (black dashed line) as fit to the experimental data (purple circles). Inset shows a zoom of the low-q region of the curves. The χ^2^ goodness-of-fit between the calculated and experimental curves is 16.5. e) The inter-atomic distance distribution calculated explicitly from the atomic model shown in **(b)** and **(c)** (black dashed line), and from the experimental data (purple line) using GNOM. Both distributions have been scaled to unity area.

### SAXS of resolvase – site II complexes

Small angle x-ray scattering (SAXS) was used to examine the conformation of Tn3 resolvase – site II complexes. SAXS was coupled with in-line size exclusion chromatography (SEC-SAXS) in order to capture scattering only from fully formed complexes (Figure 5 supplement 1a). An *ab initio* dummy atom reconstruction generated from scattering data sampled from the lowest retention-time size exclusion peak gave an envelope remarkably similar to the site II complex model described above (Figure 5b,c), despite the crudeness of the underlying assumptions required for such an analysis (e.g, without accounting for flexibility or differences in the average electron density of protein and DNA). A theoretical SAXS curve calculated from the model agrees with the actual SAXS data with a chi2 of 16.5 (Figure 5 d,e). Most of the deviation between the model and experimental data occurs at very low q, a region of the SAXS curve corresponding to low resolution shape information, such as the radius of gyration (Rg) of the scattering particle. The SAXS curve predicts a particle with a Rg of 48.9 Å, whereas the corresponding model atoms form a more compact particle with a Rg of 40.8 Å. This difference in compactness is also evident from comparison of inter-atomic distance distributions extrapolated from the SAXS data with those calculated from the model (Figure 5 d,e). A simple explanation for this discrepancy would be that the site II complex DNA is less bent and/or less stably bent in isolation than when in the context of the synaptosome. Overall, the SAXS data strongly support the primary features of the site II complex model.

### A structure-based synaptosome model

The full synaptosome was modeled using the structure and model described above for sites III and II, respectively, and the crystal structure of a site I – bound γδ resolvase tetramer which was determined by the Steitz lab using a constitutively active mutant (Figure 6)(Kamtekar et al., 2006; Li et al., 2005). To sketch out a protein scaffold, additional resolvase dimers were docked to the site IR-bound subunits, using only *R* interfaces as seen in the crystal structures, giving a V-shaped structure with site I at its base. The bound DNAs could be plausibly connected only if the middle pair of dimers was assigned to site III and the outer ones to site II. To complete the model, we based the 22-bp site I – II linker on a segment of nucleosomal DNA. The bending in the site I-II linker and at the A-tract – site IIL junction is not stabilized by directly bound protein, but agree with enhanced DNA cleavages seen by DNase I footprinting of synaptosomes (see below and Figure 6). There are small gaps between modeled DNA segments that could be sealed by small relative motions of the proteins or distortions of the DNAs.

**Figure 6.**
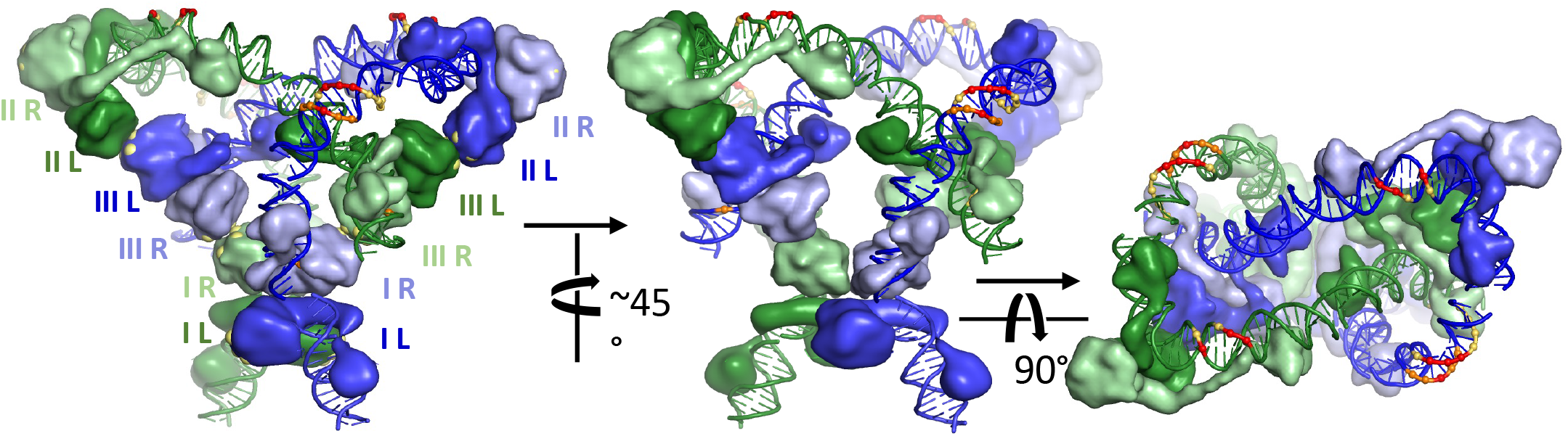
Model for the Tn3 Synaptosome Three views of the model are shown. Proteins are shown as smoothed surfaces, and are colored according to the DNA segment they are bound to, with those bound to the right half of each site in a lighter shade than those bound to the left. In the left panel, each protein is labeled according to which half site it is bound to. Also in the left panel, the alpha-carbons of key *R* interface residues (R2, R32, K54 & E56) are shown as large yellow balls poking out of the protein surface. Colored spheres and DNA backbone segments mark positions of enhanced cleavage in synaptosome footprinting experiments (see Figure 6 Supplement 1), with red marking the strongest enhancements and yellow the weakest.

Topological studies established that the synaptosome traps 3 negative plectonemic nodes between the two substrate duplexes: that is, 3 right-handed duplex-over-duplex crossings. The model is in full agreement with those studies. It shows how these nodes are trapped by the phasing of the bends introduced at sites II and III, combined with the *R*-interface contacts between the dimers bound at sites II and III of different DNA duplexes. The model is also in agreement with previous data showing that even in the absence of site I, sites II and III can support formation of a synaptic complex that traps 3 negative supercoiling nodes (e.g. (Kilbride et al., 2006)).

### Footprinting of the synaptosome

We used DNase I footprinting to interrogate the structure and accessibility of the DNA within the synaptosome. Synaptosomes were formed on specifically ^32^P-labeled supercoiled plasmid substrates, then crosslinked with glutaraldehyde before nicking with DNase I and analysis of the labelled nicked DNA by denaturing polyacrylamide gel electrophoresis.

DNase I footprinting of Tn*3*/γδ resolvase binding to *res*, and to the isolated binding site II of *res*, has been published (Grindley et al., 1982; Kitts et al., 1983)(Blake et al., 1995). The footprints of the synaptic complexes are strikingly similar to these earlier non-synapse footprints (suggesting similar accessibility of DNase I to the DNA minor groove), except for the presence of short regions of sequence where cleavages are significantly enhanced (marked on the structural model in Figure 6, and with red spots in Figure 6 supplement 1). These suggest that incorporation into the synaptosome increases DNA bending in two locations that are not in close contact with protein: within the right side of the long site II spacer region, and between sites I and II.

### Targeting *res* half-sites *in vivo*

We previously used hybrid Tn3 and *Bartonella* (Bart) resolvases and synthetic *res* sites to investigate dimer-dimer interactions in the Tn3 synaptosome during recombination (Rowland et al., 2020). The Tn3 and Bart resolvase DBDs differ in their DNA binding selectivity, and can therefore be used to target chosen catalytic domains (WT or mutant; Tn3 or Bart) to specific positions (site I, II or III) within each recombining *res*. The Tn3 and Bart resolvases also differ in their *R* interfaces, which are mutually incompatible, but crucially, each catalytic domain (Tn3 or Bart) can be switched to the other specificity by ‘transplanting’ a small surface patch of *R* interface residues from one catalytic domain to the other.

In our published work (Rowland et al., 2020) resolvases of specific types were targeted to full (dimer-binding) sites within the synaptosome. However, to test our new synaptosome model thoroughly, we needed to target individual subunits of a particular type to specific half-sites. We first designed resolution substrates to target mutant catalytic domains to both copies of a given half-site (i.e. one in each *res*) (Figure 7 and Figure 7 supplement 1). In one set of six substrates, all the half-sites bind the Tn3 resolvase DBD except for one pair (for example, site IL) that is recognized by the Bart DBD, and in a second set of six substrates, all the half-sites are recognized by Bart DBDs except one ‘Tn3’ pair. In Figure 7, results from substrates with one pair of complete Tn3 or Bart sites are included for comparison. Recombination was assayed *in vivo* as described (Rowland et al., 2020). Briefly, the substrate plasmid is transformed into *E.coli* cells expressing two resolvases, one with a Tn3 DBD, the other with a Bart DBD. Recombination (resolution) deletes a marker gene from the substrate. Transformants give ‘white’ colonies on indicator plates when resolution is efficient (shown as ‘+’ in Figure 7). When resolution is slow, colonies are red, and % resolution is then determined by analysing plasmid DNA after further growth.

**Figure 7.**
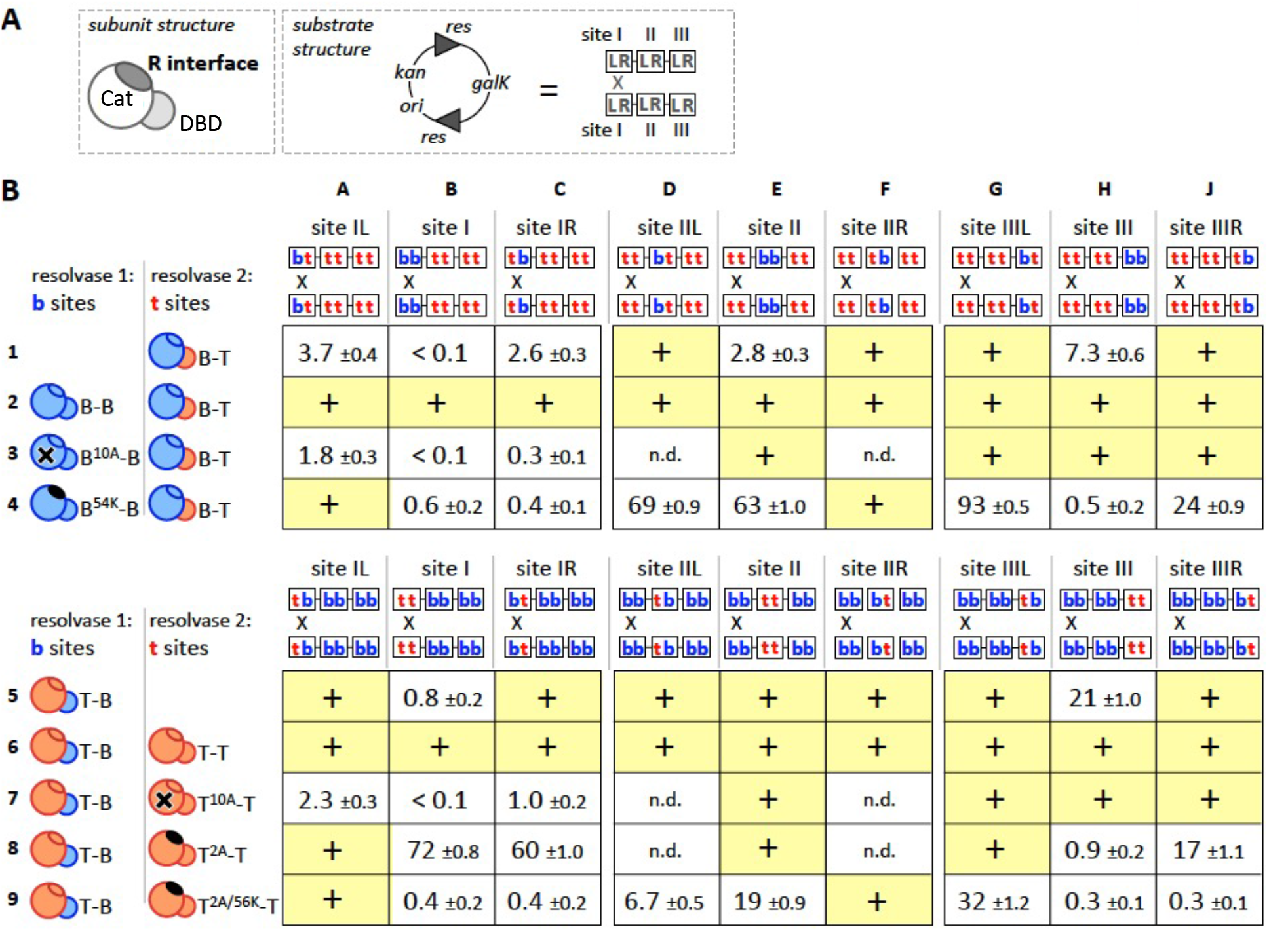
The *R* interface is required at four of six half-sites for recombination *in vivo* a) Diagrams of a resolvase subunit and a resolution substrate. The catalytic domain (Cat), DBD and *R* interface residues are labelled. All substrate plasmids have two recombination sites (*res*) in direct repeat, flanking a marker gene (*galK*). A *res* site has three dimer binding sites (I, II, III), each with left and right half-sites (L, R). The recombination crossover point (‘X’) is shown. See Figure 7 – supplement 1 for DNA sequences. b) Recombination of hybrid sites. Protein cartoons are colour coded *red* for Tn3 and *blue* for Bart. Positions of Tn3 (**t**) and Bart (**b**) half-sites in the *res* sites are shown above each column. In all rows, ‘Resolvase 1’ has a Bart DBD whereas ‘resolvase 2’ has a Tn3 DBD. In rows 1-4, all resolvases have Bart catalytic domains; in rows 5-9, all resolvases have Tn3 catalytic domains. Superscripts and symbols indicate catalytic domain mutations: S10A (‘**10A**’; **X** on catalytic domain) abolishes catalytic activity, and the others R54K (‘**54K**’), R2A (‘**2A**’) and R2A/E56K (‘**2A/56K**’) have a defective *R* interface (solid black ovals). Efficient recombination of the substrate (>90%) on the time-scale of colony growth gives ‘white’ colonies on indicator plates, shown as ‘+’ with a yellow highlight. Slower recombination results in red colonies; for these assays % recombination was determined after further growth of cells, and is shown with the standard error linked to the counting statistics (see Materials and Methods). n.d.: not done.

First we examined the effectiveness of targeting on the substrates shown in Figure 7. Recombination proceeded efficiently when the same WT catalytic domain was targeted to all twelve half-sites in the plasmid substrate (Figure 7b, rows 2 and 6). Recombination was also efficient when a catalytically inactive S10A domain was targeted to half-sites of site II or site III, but not when it was targeted to one or both half-sites of each site I (rows 3, 7). These data, showing a requirement for catalytically active subunits at site I, are consistent with the expected targeting. Targeting is most effective when the two different DBDs compete for their cognate binding sites rather than when only one type of DNA binding domain is present – e.g. compare the reactions in column C rows 3 and 7 (in which two resolvases were in competition) vs. rows 1 and 5 (in which any activity seen must reflect some resolvase subunits binding non-cognate half-sites, presumably through cooperative interactions with others). However, in some cases such as reactions A1 and C1 a single non-cognate motif can be sufficient to almost entirely block productive dimer binding.

To examine the effect of *R* interface defects at each half-site, we used Tn3 or Bart catalytic domains with *R* interface mutations (Bart R54K, Tn3 R2A or R2A/E56K) that block recombination when targeted to all sites in *res* (< 0.1% resolution in our assay; (Rowland et al., 2020)). Targeting Bart R54K domains to individual half-sites gave strong inhibition at half-sites IR and IIIR, weaker effects at IIL and IIIL, and no detectable inhibition at IL or IIR (Figure 7b, row 4). Targeting with Tn3 R2A/E56K (double mutant) gave essentially the same hierarchy of inhibitory effects at these half-sites (Figure 7b, row 9). The Tn3 R2A domain (single mutant) gave weaker inhibition at sites IR and IIIR, and no detectable inhibition at site II or IIIL (Figure 7b, row 9). This is consistent with earlier evidence that R2A at site I fails to block all recombination (Grindley, 1993; Hughes et al., 1990; Murley and Grindley, 1998), and suggests that the Tn3 *R* interface retains partial functionality with this single mutation in one partner. As reported previously (Rowland et al., 2020), targeting the Bart R54K domain to both halves of site I or site III (i.e. as dimers) strongly inhibited recombination, but the effect was notably weaker at site II (Figure 7b; reactions B4, E4, H4).

In summary, efficient recombination requires *R* interface proficiency at four of the half-sites (IR, IIL, and both halves of site III), but not at the other two (IL and IIR), consistent with the synaptosome model (Figure 7a). Notably, recombination is more sensitive to *R* interface mutations at sites IR and IIIR than it is at sites IIL and IIIL.

### Evidence for IR-IIIR and IIL-IIIL *R* interactions in *trans*

In the experiments described above, we used catalytic domains with *defective R* interfaces to identify the subunits making critical *R* interactions. To investigate which pairs of subunits interact within the synaptosome, we needed to use two resolvases with *functional R* residues. We aimed to assemble synaptosomes containing both Tn3-type and Bart-type *R* interfaces, reasoning that efficient recombination would require subunits positioned such that all connected dimers have matching *R* residues. The following experiments ask in particular which site III-bound subunit interacts (via the *R* interface) with a site IR-bound subunit and which interacts with a IIL-bound subunit, and whether those interactions are in *cis* or in *trans* (i.e. between subunits bound to the same *res* site or to partner *res* sites).

We showed previously that *R* residues can be ‘transplanted’ between Tn3 and Bart resolvases, to confer the *R* interface specificity of the donor, and that catalytic domains with matching *R* residues interact most productively (Rowland et al., 2020). Thus to provide Tn3-type *R* residues, we used either a WT Tn3 catalytic domain or a Bart catalytic domain with Tn3-type *R* residues (B^T^), and to provide Bart-type *R* residues we used a WT Bart catalytic domain or a Tn3 catalytic domain with Bart-type *R* residues (T^B^).

We used a set of eight substrates containing *res* sites with different arrangements of Tn3- and Bart-type half-sites (that is, targeted by Tn3 or Bart DBDs; Figure 8a). All substrates have one Tn3 site I and one Bart site I, so different types of catalytic domain can be targeted to each. Because only site III required *R* interface proficiency at both half sites, oriented heterodimers were targeted exclusively to that site (substrates Figure 8a; substrates P through S). The new synaptosome model predicts that the subunits at sites IR and IIIR pair *in trans;* likewise the subunits at IIL and IIIL (Figure 8b). Figure 8c shows how the test substrates are configured: subunits targeted to IR and IIIR (or to IIL and IIIL) will match either *in trans*, or *in cis*.

**Figure 8.**
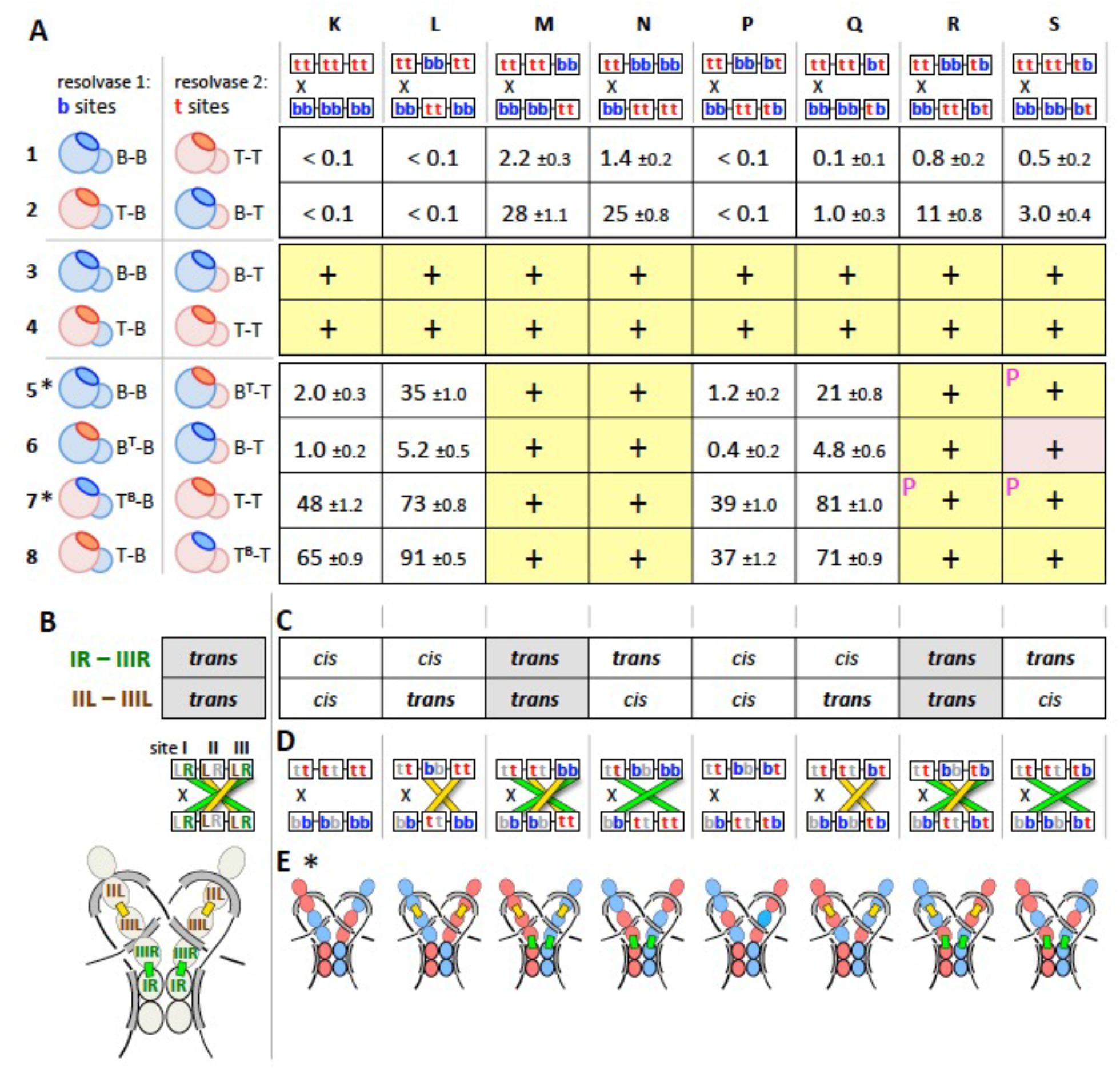
Evidence for *R* interfaces *in trans* between sites IR and IIIR, and sites IIL and IIIL. a) Recombination of hybrid substrates. Substrates and resolvases used are diagrammed as in Figure 7, but with the *R* residue patches highlighted in darker shades indicating their specificity. ‘Swapped’ *R* interfaces are also denoted with superscripts (T^B^: Tn3 catalytic domain with Bart *R* interface residues; B^T^: Bart catalytic domain with Tn3 *R* interface; see text for details). All boxes for which recombination was efficient enough to detect by a change in colony color are marked with a ‘+’. These are further differentiated as follows: a yellow background indicates the most efficient recombination (white colonies); yellow with a P indicates almost white (pale pink) colonies, and a pink background indicates pink colonies (and therefore partial resolution). Slower recombination results in unchanged (red) colony color; for these assays % recombination was determined after further growth of cells, and is shown with the standard error linked to the counting statistics (see Experimental Procedures). b) Predicted subunit contacts in the new synaptosome model. Top: tabulation of the predicted interactions for sites IR and IIIR (*green*) and for sites IIL and IIIL (*brown*/*yellow*) (*trans* meaning between the two paired *res* sites). Middle: recombination substrates are diagrammed as in Figure 7a, with colored bars depicting the predicted contacts between the two partner sites (green, IR-IIIR; yellow, IIL-IIIL). Bottom; a cartoon of the synaptosome model showing a more realistic view of these connections (see also Figures 6 and 10). c) Tabulation of cognate *R* interactions between sites IR and IIIR and sites IIL and IIIL that can be made on each experimental substrate. Only two substrates are configured so that both of these pairs of half-sites will match *in trans* (grey highlight) - the arrangement that corresponds to the model (see (b)). d) Diagram of the *trans* interactions that can be formed for each substrate. Bars connecting interacting sites are colored as in (b). Cartoons of the synaptosome model. The *R* interactions that are both predicted by the model and allowed by each substrate are highlighted as green and yellow bars. Subunits are colored according to the *R* interface residues of the subunit targeted to each half-site (red, Tn3; blue, Bart) in rows 5 and 7 of part (a); (indicated by *).

The first four rows of Figure 8 demonstrate the importance of cognate interactions among catalytic domains, regardless of which DBD is attached. All eight substrates recombined efficiently when the twelve half-sites were targeted by the same catalytic domain (Figure 8a, rows 3, 4). In contrast, these substrates recombined inefficiently when the pairs of resolvases used had different catalytic domains (rows 1, 2). This is as expected (even for substrates allowing compatible *R* interfaces), because a functional catalytic tetramer cannot be assembled at a pair of site Is from two Tn3 and two Bart catalytic domains, as seen previously (Rowland et al., 2020). In rows 5-8, the catalytic domains are the same within each pair of resolvases except that their *R* residues are varied.

When the resolvases are of the same catalytic domain type, but have non-matching *R* residues (rows 5-8), efficient recombination occurred in the four substrates where the IR and IIIR motifs match *in trans* (columns M, N, R, S). These data argue strongly that *R* interactions *in trans* between the catalytic domains at sites IR and IIIR are a key determinant of recombination efficiency. This is consistent with our evidence that *R* interface proficiency is most critical at sites IR and IIIR (Figure 7b), and with the synaptosome model (Figure 6).

Deciphering the *R* interactions of site IIL was complicated by the fact that *R-deficient* dimers targeted to site II have a relatively weak (Figure 7b: E4, E9), or, in one case, undetectable (E8) effect. When the resolvases at sites IR and IIIR match *in trans*, no differential effect of a match or mismatch between sites IIL and IIIL was detectable: recombination was efficient in all cases (Figure 8a: rows 5-8, columns M, N, R, S; the assay ‘saturates’, and may not detect small changes in absolute rate). This result suggests that mismatched *R-proficient* dimers at site II do not strongly inhibit recombination (assuming site I and site III are matched), consistent with our findings in Figure 7b. However, when the resolvases at sites IR and IIIR do *not* match *in trans*, we consistently see faster recombination when the resolvases targeted to sites IIL and IIIL match *in trans*, rather than *in cis* (Figure 8a, rows 5-8; columns K, L, P, Q). The differences are clearest for the B-B + B^T^-T resolvase pair (row 5); for the other pairs the differences may be reduced by residual compatibility of the Tn3 and Bart-type *R* residues, or by partial binding of DBDs at non-cognate motifs. All available data clearly indicate that the *R* residues in subunits IIL and IIIL are important for recombination (Figure 7b), and for accessory site synapsis (Murley and Grindley, 1998), and that no other interface has a bridging role at sites II and III (Rowland et al., 2020). We thus deduce that subunits IIL and IIIL are linked *in trans*, as specified by the new synapse model (Figure 6).

## Discussion

The new model for the Tn3 resolvase synaptosome presented here agrees with a large body of historical data for the Tn3 and γδ systems (reviewed in (Rice, 2015; Stark, 2014)) and also the related Tn21 resolvase system (Soultanas et al., 1995; Soultanas and Halford, 1995). It differs from previous models in proposing a less compact protein core, which is consistent with our data showing that the only functionally important inter-dimer interactions formed by the site II- and III-bound subunits are those involving the *R* interface. In contrast, the earlier models proposed more extensive contacts among the resolvase subunits (see below).

The delicate appearance of our synaptosome model probably reflects its natural function. The synaptosome needs to be robust enough to bring together the two *res* sites in the cointegrate product of replicative transposition, which are separated by the length of the transposon (~5 kbp) and the donor/target replicons (Figure 1a). However, its stability may be finely tuned so as to favour formation of the resolution product. Experiments *in vitro* show that strand exchange by a rotational mechanism terminates predominantly after only a single 180° rotation of the two site IL-bound subunits (yielding a 2-noded catenane resolution product; Figure 1c), and our current hypothesis is that this selectivity is due to rapid dissociation of the synaptosome. After one round of 180° rotation, negative supercoiling no longer favours wrapping of the DNA as in our model, and complete dissociation of the two (recombinant) *res* sites would release substantial free energy. (reviewed in (Rice, 2015; Stark, 2014)). A loose flexible synaptosome architecture might kinetically favour this dissociation.

The new model differs from previous ones in the details of the inter-dimer interactions. In the new model, each resolvase dimer at site III contacts dimers at sites I and II *in trans (ie*. bound to the partner *res*), using *R* interfaces between subunits at sites IR and IIIR, and at sites IIL and IIIL (Figures 6 and 8). This contrasts with three earlier models where the *R* interface connects dimers *in cis* at sites II and III (Murley and Grindley, 1998; Rice and Steitz, 1994b; Sarkis et al., 2001), and models where it connects dimers *in cis* at site I and site III (or site II in the Sin system), as in a structure-based model of the Sin synapse (Mouw et al., 2008), and a similar model for the Tn3 resolvase synapse (Rowland et al., 2009). The data presented here (Figures 7 and 8) strongly support the specific set of dimer contacts seen in the new model, and rule out other suggested arrangements (see Figure 8 Supplement 1). The *trans*contacts proposed in the new model are also consistent with earlier observations that a pair of truncated *res* sites comprising just binding sites II and III can nevertheless form an interwrapped synaptosome (Kilbride et al., 1999; Watson et al., 1996).

It has long been argued that a primary reason for the interwrapped DNA architecture of the Tn3 synaptosome is to implement a mechanism called topological selectivity. Recombination by Tn3 resolvase, *in vivo* and *in vitro*, has the remarkable property that only pairs of *res* sites in direct repeat (head to tail) relative orientation in a supercoiled circular plasmid molecule recombine efficiently. There is very strong selectivity against recombination of *res* sites in inverted repeat orientation, or on separate DNA molecules, or in non-supercoiled DNA molecules (rates are typically ~10^3^-fold or more slower) (Brown et al., 2002; Watson et al., 1996). It is proposed that formation of an interwrapped synaptosome trapping 3 negative supercoil nodes is energetically favourable in a standard supercoiled plasmid substrate with directly repeated *res* sites, but is highly disfavoured for other arrangements of sites due to distortions of the substrate DNA that would be necessary (Boocock and Stark, 1995). Our new synaptosome model is entirely consistent with this selectivity mechanism, as well as the proposed mechanism for selective formation of 2-noded catenane recombinant product (see above).

How does synaptosome formation induce a conformational change in the site I – bound proteins from the catalytically inactive dimer to the catalytically active tetramer? A portion of the free energy released by the formation of multiple favorable protein-protein and protein-DNA contacts in the synaptosome must be used not only to hold together the two otherwise-distant *res* DNA segments but also to drive this conformational change in the site I-bound proteins. In part that free energy may be used to fight entropy by aligning the two site I dimers properly and holding them in very close proximity. We previously demonstrated that mass action (through high solution concentration) can drive catalytic activation of Sin resolvase, albeit weakly (Mouw et al., 2010). However, the synaptosome architecture might also directly strain the site I-bound subunits so as to favour the active conformation. As diagrammed in Figure 9, fitting the site IR subunit of an inactive WT resolvase dimer into the synaptosome model requires moving the E helix and the DNA bound to it relative to the catalytic core domain (and its *R* interface) in a way that naturally pulls it towards the active conformation. The location of the *R* interface patch roughly opposite the DNA-binding end of helix E may maximize the leverage applied by the synaptosome.

**Figure 9.**
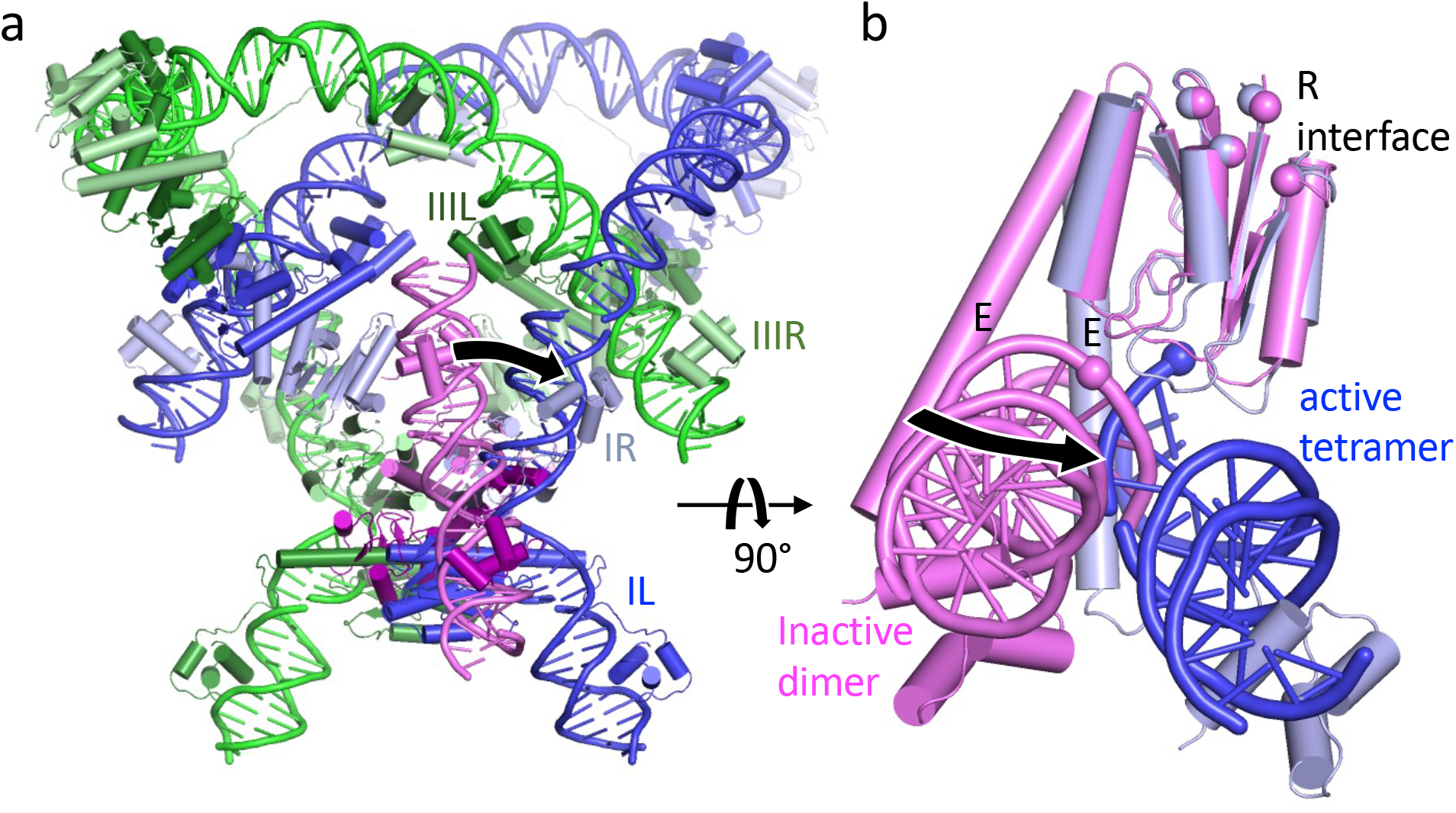
The dual constraints of the DNA path and the *R* interface may aid activation a) Reference view. The Tn3/ γδ synaptosome model, with the site I – bound WT γδ resolvase dimer (1GDT; pink & purple (Yang and Steitz, 1995)) superimposed onto the site I – bound tetramer, using the catalytic domains of the site IR-bound (light pink and light blue) subunits as guides. Arrow shows motion of the DNA required for the *R* interface to lock together the site IR and site IIIR bound subunits as required for synaptosome formation. b) Closeup view, rotated 90° about a horizontal axis and showing only the superimposed subunits. Cα atoms of residues R2, R32, K54, and E56 are shown as spheres to mark the *R* interface where this subunit interacts with that bound to site IIIR (light green in part a). The arrow shows how, if the *R* interface has already locked the catalytic domain into place within the synaptosome, adjusting the pink subunit to minimize strain in the DNA shifts the E helix from the dimer conformation towards the tetramer conformation (relative to the catalytic domain). The scissile phosphates are also marked with a sphere.

Figure 10 compares the new model for the Tn3 synaptosome with our previous model for the Sin synaptosome (Mouw et al., 2008). Sin and its close relatives also function as resolvases but are typically encoded by large theta-replicating *S. aureus* plasmids rather than replicative transposons (LeBard et al., 2008). Like Tn3, the Sin synaptosome traps 3 negative supercoils – a topology that is likely conserved for its utility in channeling recombination to produce only intramolecular resolution products (see above). However, the accessory sites for Sin are quite different (Figure 1b), and it uses a host DNA-bending protein in place of Tn3’s site II (Rowland et al., 2005). Nevertheless, Sin synaptosome formation also requires *R* interface interactions between the site I – bound dimers and the dimers bound to the opposite end of each *res* site.

**Figure 10.**
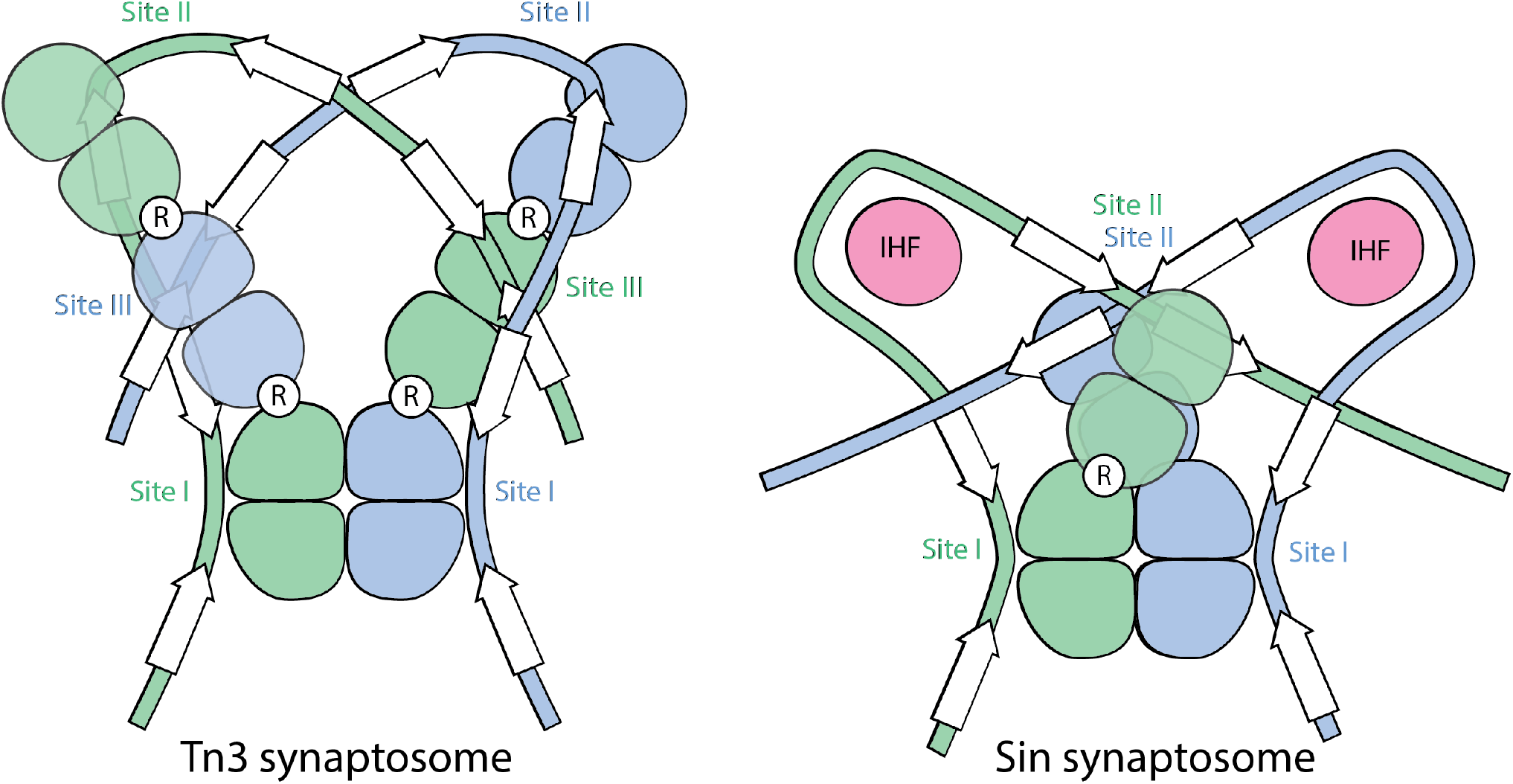
Tn3 and Sin resolvases use different synaptosome architectures to trap similar DNA topology. The cartoons show only the positions of the catalytic domains, with arrows for the DNA motifs that the DBDs recognize. *R* interfaces are marked with white circles.

The *R* interface patch of the protein is also important in the activation of Hin, a small serine recombinase that, in its natural setting, catalyzes inversion rather than resolution. In the Hin case, the crossover sites for Hin (equivalent to *res* site I) are synapsed within an “invertasome” that involves a different topology and a different protein cofactor (Johnson, 2015). In this case the *R* interface-like residues contact DNA rather than another protein (McLean et al., 2013). The repeated use of this patch of protein surface for intermolecular interactions within the synaptic complexes that activate small serine recombinases supports the idea that such interactions may be more than simply scaffolding.

Our structural data also highlight how the overall geometry of the complexes of both Tn3 and Sin resolvase dimers with their DNA accessory sites is largely determined by the spacing and orientation of three sequence motifs in the DNA: the preferred sequence for helix E docking into the minor groove, and the two specific sequences for binding the helix-turn-helix DBDs of the dimer (Figures 3 and 5). The proteins adapt to this geometry by use of a highly flexible linker between the end of helix E and the DBDs and flexibility at a kink-prone position within helix E. Interactions between the non-DNA-binding face of helix E and the catalytic domain may also help to determine the orientation of the catalytic domain dimer relative to the DNA. However, that interaction is not extensive (see Results section) and we cannot rule out that, when free in solution, the catalytic domains display some mobility about the helix E hinge. The modular nature of these proteins may have facilitated the evolution of different synaptic complexes that channel recombination differently towards the biologically relevant outcome – e.g. resolution for Tn3 and Sin resolvases, but inversion for Hin. The geometry of large serine integrase – DNA complexes is also dictated by the spacing and orientation of DNA sequence motifs but in a different manner: different dimer-binding sites have different spacings between the cognate subsites for each of the integrases’ two C-terminal DBDs (Rutherford et al., 2013).

Understanding the construction of the Tn3 synaptosome and how it controls the activity of the crossover site-bound proteins paves the way to further engineering of these systems for use as genomic tools. While serine recombinases lack the flexibility in target sequences of CRISPR-Cas systems, they do not create R-loops or leave double-strand breaks for the host to repair. Furthermore, the modular nature of small serine recombinases facilitates their engineering to alter target specificity: for example, constitutively active mutants of their catalytic domains can be fused to other DNA binding domains such as zinc fingers and even Cas9 (Mercer et al., 2012; Proudfoot et al., 2011; Zhang et al., 2021). However, because those chimeras are constitutively active, they will synapse any pair of sites regardless of topological context, and will catalyze (usually undesirable) further rounds of recombination between the product sites. Other commonly used recombinases (for example, the tyrosine recombinases Cre and Flp) behave similarly (Grindley et al., 2006)(Van Duyne, 2015). The large serine integrases use an additional protein to ensure a single round of recombination and to toggle between the forward and reverse reactions (‘directionality’). The integrase alone catalyzes insertion (integration) of phage DNA into the host chromosome, but not the reverse (excision) reaction; however, binding of a small phage-encoded ‘recombination directionality factor’ protein to the integrase alters the protein-DNA complex so that it catalyzes only excision (Smith, 2015). Engineering the sequence specificity of the large serine recombinases is difficult because of the structural complexity of the protein-DNA interactions and because the cofactor protein binds to the DNA binding domain itself. What is now known about Tn3 resolvase and its controlling synaptosome should allow for further engineering to exploit this system for directing particular recombination outcomes (e.g. insertion, deletion or inversion) when and where desired.

## Materials and Methods

### Sample Preparation

Tn3 resolvase was overexpressed in Rosetta DE3 pLysS cells from plasmid pSA1121, which encodes wild-type Tn3 resolvase (GenBank: CAA23885.1) with SHHHHHH added after the natural C-terminus. pSA1121 is identical to pSA1122 (Rowland et al., 2005) except that the Sin reading frame is replaced by that for Tn3 resolvase). Cells were grown at 37°C in Luria-Bertani medium containing 50 μg/ml kanamycin and 10 ug/ml chloramphenicol to an OD_600_ of ~0.8 before 0.5 mM IPTG was added. Cells were then grown for 3 additional hours. Cells were harvested by centrifugation at 8000 rpm in a Fiberlite F10-6 x 500y rotor for 10 minutes. Cell pellets were resuspended in a lysis buffer (50 mM Tris pH 8.0 and 0.3 M NaCl), sonicated, and centrifuged at 18000 rpm in an SS-34 rotor for 1 hour. Tn3 resolvase remained in the pellet and was solubilized by resuspending the pellet in a denaturing buffer containing 0.1 M phosphate pH 8.0, 0.3 M NaCl, and 7 M urea. After resuspension, the mixture was filtered, and the filtrate was loaded on a Ni-HiTrap column (GE Healthcare). The protein was eluted by applying an imidazole gradient from 20 mM to 250 mM. Protein fractions were pooled, and dialyzed into a buffer containing 50 mM Tris pH 8.2 and 7 M urea. The sample was then loaded on a Mono-S ion-exchange column (GE Healthcare). A salt gradient from 50 mM to 2 M NaCl was applied to the column to separate out the Tn3 resolvase from other impurities. Surprisingly, upon inspection of the fractions on an SDS-PAGE gel, most of the impurities and a small fraction of Tn3 resolvase bound to the column, leaving the flow-through with Tn3 resolvase protein of high purity. This highly pure Tn3 resolvase was refolded by dialyzing at 4°C into a buffer containing 25 mM Tris pH 7.5, 2 M NaCl, and 15% glycerol. The dialysis was done four times, each in a buffer volume 5 times that of the Tn3 sample. The protein was further dialyzed into 25 mM Tris pH 7.5, 0.4 M ammonium sulfate, 0.5 mM EDTA, and 20% glycerol and concentrated to ~35 mg/ml.

Unmodified DNA oligonucleotides were obtained from Integrated DNA Technologies. DNA oligonucleotides that were labeled with 5-bromo-dU for experimental phasing purposes were ordered from Keck Oligonucleotide Synthesis Facility (Yale University). Each DNA duplex used in crystallization was prepared by mixing two complementary single strands in equimolar amounts, heating at 80°C for 20 minutes, and annealing by slow cooling to room temperature. The final concentration of the duplex DNA was 1 mM in a TE-buffer (Tris-HCl pH 8.0-EDTA) containing 50 mM NaCl. Form I crystals were obtained using the following oligonucleotides: 5’-TCG**TGTCTGA**TATTCGATTTA**AGGTACA**TT and 5’-<u>A</u>AA**TGTACCT**TAAATCGAATA**TCAGACA**CG, while Form II crystals were obtained using the following: 5’-<u>AT</u>**TGTCTGA**TATTCGATTTA**AGGTACA** and 5’-<u>AT</u>**TGTACCT**TAAATCGAATA**TCAGACA**, where bold nucleotides mark the DBD binding site motifs and underlined nucleotides were changed from the native *res* site to facilitate crystal packing.

### Crystallization and Data collection

All complexes were formed by mixing Tn3 resolvase with DNA in a 2:1.25 molar ratio. They were incubated for at least 30 minutes at room temperature prior to setting up hanging drops for crystallization. The Form I crystals were obtained from a 6 mg/ml complex (in 25 mM Tris pH 7.5 and 0.18 M ammonium sulfate) mixed in a 1:1 ratio with well solutions containing 19-21% PEG3350 and 0.2 M sodium malonate pH 7.0. Microseeding was used to acquire larger single crystals suitable for data collection. To obtain the microseeds, clusters of crystals were finely crushed, stabilized in the well solutions, and serially diluted. From each dilution, 1 μl of microseeds were added to 1 μl of the complex to form the hanging drop, which was then suspended over the precipitant solution. Drops were incubated at 19 °C for crystal growth. Tantalum bromide derivatives for Form I were obtained by soaking the crystals with 22% PEG3350, 0.2 M sodium malonate pH 7.0, 0.18 M ammonium sulfate, and 0.4 mM tantalum bromide cluster, [Ta_6_Br_12_]^2+^ * 2 Br– (Jena Bioscience) for 1 to 5 days. Bromine derivatives were obtained by using brominated DNA where all but the terminal thymines were replaced with 5-bromo-dU. Native crystals were frozen in liquid nitrogen under conditions that mimicked the drop supplemented with a 20% PEG400/10% glycerol cryoprotectant mix. Derivative crystals were frozen in a similar manner but with 20% glycerol.

Form II crystals were obtained from a 3 mg/ml complex (in 25 mM Tris pH 7.5 and 0.14 M ammonium sulfate) mixed in a 1:1 ratio with well solutions containing 16% PEG3350 and 0.2 M ammonium fluoride. Like the Form I crystals, Form II crystals were also derivatized with Ta_6_Br_12_. They were soaked for 5 days in 16% PEG3350, 0.2 M ammonium fluoride, 0.14 M ammonium sulfate, and 0.4 mM Ta_6_Br_12_ before being frozen with liquid nitrogen in a similar solution but with 20% glycerol. The native crystals were frozen in much the same way but without the tantalum cluster and using 20% ethylene glycol as cryoprotectant.

All data sets were collected at SBC-CAT 19-ID beamline. X-ray data from the native crystals were integrated and scaled with HKL2000 suite and the others with HKL3000 (Otwinowski and Minor, 1997). A summary of the data collection statistics is shown in Tables 1 and 2.

### Structure Determination and Refinement

The Form I native data set extends to about 3.4 Å resolution. Phases were determined by MIRAS (Multiple Isomorphous Replacement and Anomalous Scattering) using two derivatives, Ta_6_Br_12_ and Br. Initially, a single Ta_6_Br_12_ cluster was identified by direct methods (SHELXD) as well as by anomalous difference Patterson methods implemented in SOLVE (Sheldrick, 2008; Terwilliger and Berendzen, 1999). This cluster was consistent with the anomalous difference Patterson maps generated in CNS from Ta_6_Br_12_ data sets where crystals were soaked for 3 and 5 days (Brunger, 2007). SIRAS phases generated from the position of this one Ta_6_Br_12_ cluster were then utilized to locate the positions of the Br atoms in the brominated DNA data set. Inspection of these Br sites suggested that there is only one DNA molecule, and hence one Tn3 resolvase dimer-Site III DNA complex, in the asymmetric unit, and that the solvent content is therefore 66% solvent. Analyzing the peak heights of the Br sites confirmed resolution of the space group ambiguity in favor of P4_3_2_1_2 rather than P4_1_2_1_2. Using SIRAS phases from the Br atoms allowed us to find two more Ta_6_Br_12_ clusters. Ultimately, the phases for the Form I structure were calculated using anomalous and isomorphous signals from 3 Ta clusters and 12 Br atoms. Phases were improved by density modification using Parrot in CCP4 (see Figure 2 supplement 1 for experimentally-phased electron density map) (Winn et al., 2011). The model of our Form I structure was built in COOT using previously determined structures from apo γδ resolvase and its complexes with Site I-DNA (PDB ids 2rsl and 1gdt) (Emsley and Cowtan, 2004; Rice and Steitz, 1994a; Yang and Steitz, 1995). Helix E had to be adjusted considerably in our structure because it is very different from that of previous γδ and Sin resolvase structures. Confidence in our model comes from the fact that Tn3 and γδ resolvases share >90% sequence identity in their catalytic domains and ~48% in their DNA-binding domains. In addition, the bromine peaks guided the building of our DNA model, and clearly confirmed the directionality of Tn3 binding. Despite the fact that the protein is a homodimer, the DNA sequence and the complex formed with it are highly asymmetric, and there were no bromine peaks suggesting twofold disorder in the DNA’s orientation. The structure was refined using PHENIX (Adams et al., 2010). During the course of refinement and manual rebuilding, Ramachandran restraints, secondary restraints, H-bonding restraints on the DNA, and NCS averaging on both the catalytic and DNA-binding domains were executed. The final refinement used data to 3.4 Å, NCS restraints, Ramachandran restraints, grouped B-factors (2 per residue) and 9 TLS (translation/libration/screw) groups. The final R_work_ and R_free_ were 25.1% and 30.0%, respectively.

The Form II native data extend anisotropically to approximately 4.0/4.5/4.0 Å resolution. Initial stages of the Form II structure determination involved molecular replacement (MR) using the Tn3 resolvase DNA-binding domain bound to its 12-mer binding site (DBD-DNA) as search model. The CCP4 program, Phaser, found 4 solutions, which showed convincing crystal packing arrangements in that the spacing between two DBD-DNA’s corresponded to a single base pair, and the crystal contacts were in agreement with a tail-to-tail packing of the DNA ends. This information indicated that there are 2 Tn3 resolvase-Site III DNA complexes in the asymmetric unit. More MR methods using MOLREP were then employed to find the four catalytic domains. When only one catalytic domain was used as the search model, only two of the four could be located. However, when the catalytic dimer from the Tn3 Form I structure was used, all domains were accounted for, two of which overlap with the other two catalytic domains found in the previously described MOLREP run. To verify if these catalytic domains were positioned correctly, we utilized a low-resolution Ta_6_Br_12_ data set. Model phases generated from a model that did not include the catalytic domains were used in an isomorphous difference Fourier method to find Ta clusters. Although only one Ta cluster was found, it was enough to obtain unbiased electron density for the catalytic domains by employing SigmaA to combine the Ta_6_Br_12_-SIR phases with the model phases and using density modification in Parrot to further improve the phases. The model was later rebuilt starting from the form I model and adjusting for the slightly different bend in the DNA in form II. The Form II structure was refined in PHENIX using the same restraints and strategies of B-factor handling as were used for Form I. The final model was refined to R_work_ = 20.7% and R_free_ = 25.0%. In all 3 independent determinations of the dimer-DNA complex, the catalytic domain that is not in direct contact with the DNA was rather poorly ordered.

### Synaptosome modeling

Modeling was carried out manually using PyMOL (the PyMOL Molecular Graphics System, Schrodinger, LLC) and subsets of multiple coordinate files. One protein-bound *res* site (~120 bp and 3 resolvase dimers) was modeled explicitly, and the coordinates were assigned a fake space group of P 2 1 1 and placed such that crystallographic symmetry generates the partner *res* site. Coordinates are included as supplementary material.

The tetramer bound to site I was based on 1zr4, a structure of a constitutively active γδ resolvase mutant covalently linked to cleaved site I DNAs (Li et al., 2005). The non-crystallographic axis relating the two site I DNAs was aligned to the x axis (y=z=0). The half of this complex comprising subunits B and D was initially used, although later subunit A (and its bound half-site DNA) were superimposed on subunit D and those coordinates were used instead of subunit D’s for site I-right. Additional Tn3 resolvase dimers were placed such that they recreate the regulatory dimer-dimer interface (using pairs of Tn3 dimers as shown in Figure 4a as guides). The relatively smoothly curved linker DNA to connect sites I and II was taken from a nucleosome structure, PDB id 1eqz.

Modeling of the site II-DNA complex was based on the site III-bound crystal structure, with initial coordinates for the 9-bp insertion taken from the crystal structure of an A-tract-containing DNA, 1bdn (DiGabriele et al., 1989). Coot was used to re-optimize local geometry after merging DNA files, after unfolding the C-terminal segment of one subunit’s E helix and after adjusting DNA bending angles, which was done manually in PyMOL.

In the deposited model, the DNA sequence for site I is that of the γδ resolvase structure it was taken from, the sequence for the site I – II linker is that of the original chicken nucleosomal DNA, and that of sites II and III is that of the Tn3 *res* site.

The Sin synaptosome model presented here is based on that described earlier (Mouw et al., 2008). We updated it by replacing the γδ resolvase tetramer at site I with a model based on an activated Sin tetramer and the DNA – DBD complexes from the left site of the Sin – site II complex (Keenholtz et al., 2011). The coordinates of one protein-bound *res* site (included as supplementary material) are centered on the z axis and assigned the fake space group P 1 1 2 such that crystallographic symmetry generates the partner site.

### SAXS

SAXS data were collected at the BioCAT beamline at the Advanced Photon Source (APS) at Argonne National Laboratory. The Site II complex sample for SAXS was prepared by mixing wild-type Tn3 resolvase with site II DNA in a 2.2:1 molar ratio to a final concentration of 3 mg/mL in a buffer consisting of 25 mM Tris pH 7.5, 150 mM NaCl, 10 mM MgCl_2_ and 1% glycerol. This was incubated for at least 30 minutes on ice prior to the experiment, then passed through a 0.22 μm centrifugal filter immediately prior to injection. 500 μL of this sample was injected onto a Superdex 200 10/300 GL column which had been equilibrated in the same buffer. This chromatography was performed on an Akta Pure FPLC system running at 0.45 mL/min, the output of which was directly connected to the beamline sample cuvette. SAXS data were collected as 1.09 s exposures which were taken approximately every 2.5 s beginning at approximately 5.5 mL post injection. An overlay of the UV and the total scattering measured in each frame is given in Figure 5 – supplement 1a; as the delay volume between the FPLC UV detector and SAXS sample cuvette was not precisely known, the alignment shown is approximate.

For data reduction, the data were truncated to the range 0.01 < q < 0.2 Å^-1^. 50 frames from the beginning of the column run, prior to the column void volume and any eluting UV signal, were averaged to represent the buffer blank. 40 frames from the leading half of the lowest retention-time (first) UV elution peak were taken as “sample” frames. The buffer blank was subtracted from each of these, followed by scaling and averaging using DATMERGE program to generate the final SAXS data used for analysis (Petoukhov et al., 2007). Guinier (Rg) fitting and distance distribution extrapolation were performed using AUTORG and DATGNOM respectively (Petoukhov et al., 2007). The fit to the Site II complex atomic model was performed using FoXS (Schneidman-Duhovny et al., 2013).

### DNase I footprinting of crosslinked Tn*3* resolvase-*res* synaptic complexes

Synapsis substrate plasmids (3.2 kbp) contain two Tn*3 res* sites in direct repeat, separated (crossover site - crossover site) by 568 bp. pMS178 (used to obtain the top strand footprint shown in Figure 6 supplement 1) differs from pALY25 (used for the bottom strand footprint) only by the addition of a short oligonucleotide sequence at the site I end of one *res* site, introducing a site for the restriction endonuclease MluI.

Site-specifically ^32^P-labelled supercoiled plasmid substrates were prepared by a method adapted from that used by M^c^Ilwraith *et al*. (McIlwraith et al., 1997). The plasmid DNA was linearized by digestion with BamHI (pALY25) or MluI (pMS178) at unique sites adjacent to the *res* site III and site I ends respectively, and the 5’-phosphates were removed with calf intestinal phosphatase. Following extraction of proteins with phenol-chloroform and ethanol precipitation of the DNA, the 5’-ends were labelled with T4 kinase and [γ-^32^P]ATP. The ends were then phosphorylated to completion by addition of 0.1 mM unlabelled ATP. The DNA was precipitated with ethanol, then redissolved at low concentration (~3 μg ml^-1^) in a buffer containing 2 μg/ml ethidium bromide and recircularized with T4 DNA ligase. The labelled supercoiled plasmid DNA was then recovered by ethanol precipitation and purified by low-melting point agarose gel electrophoresis. A suitable activity of the labelled supercoiled plasmid was mixed with an excess of unlabelled plasmid prior to synapsis and footprinting experiments.

Synapses were stabilized by crosslinking with glutaraldehyde and fixing with sodium borohydride, essentially as described by Watson *et al*.(Watson et al., 1996). Nicking was initiated by the addition of DNase I and 10 mM MgCl_2_, and stopped by addition of excess EDTA.

Samples were then separated on a low-melting point (Seaplaque) agarose gel, and the band corresponding to nicked synapse (Watson, 1994) was excised. The control DNA (no resolvase) was treated identically, and the nicked circle DNA was excised. The recovered DNA was digested at the labelled site (with BamHI (pALY25) or MluI (pMS178)) and with EcoRV, which cuts the non-*res* end of the linearized DNA to release a very short labelled fragment. Samples were then loaded and run on ‘sequencing’ polyacrylamide gels (see for example (Rowland et al., 2002)). Sequence positions of bands on the gels were assigned by comparison with ‘G-track’ marker ladders, prepared by treating the labelled plasmid DNA (digested with BamHI or MluI and EcoRV, as above) with dimethyl sulphate and then piperidine. Gels were phosphor-imaged using a Fuji BAS instrument, and quantitated/profiled using ImageGauge software (Fuji).

### *In vivo* recombination assays

Resolvase expression plasmids, substrate plasmids, and the complementation assay for recombination are described in (Rowland et al., 2020). The four synthetic *res* sites used to construct the substrates are shown in Figure 7 supplement 1. Plasmid sequences are available on request. In Figures 7b and 8a, % recombination was determined as previously described by retransformation of plasmid DNA; the associated RMS sampling errors (SE) are given as: SE (%) = 100√(p.(1-p)/n), where w and r are the counts of white and red colonies, p = w/(w+r), and n = w+r. Quantitative comparisons discussed in the text are from assays done in parallel in two data sets in Figure 8: (1) rows 1, 2; (2) rows 5, 6, 7, 8; and four data sets in Figure 7: (1) columns D, E; (2) columns G, H, J; A1, A3, B1, B3, B4, C1, C3, C4; (3) A7, B5, B7, C7; (4) B8, B9, C8, C9 For nine representative reactions (with % recombination in the range 0.8 - 71%), assayed in triplicate in independent data sets, the standard deviation in the estimated % recombination was between 0.1% and 2.4% (Rowland et al., 2020).

### Visualizations

Structure figures were made using PyMOL (the PyMOL Molecular Graphics System, Schrodinger, LLC). DNA roll angles were measured using w3DNA2.0 (http://web.x3dna.org; (Li et al., 2019)). Roll angles from the “simple step” chart were reported.

### Data files

Crystallographic coordinates and reduced diffraction data for crystal forms I and II were deposited with the RCSB protein data bank, ID #s 5cy1 and 5cy2, respectively. The raw diffraction data were deposited with SBgrid, data sets #682 (10.15785/SBGRID/682) and 683 (10.15785/SBGRID/683), respectively.

## Supporting information

Table 1. Crystallographic Data Collection and Refinement Stastics.

Tn3 synaptosome model (one half with symmetry operators)

Tn3 synaptosome model 2nd half

Sin synaptosome model (one half with symmetry operators)

Sin synaptosome model 2nd half

Supplemental Movie - Figure 6 rotating

## Additional files

Tn3synap_twirl.mpg – movie of the Tn3 synaptosome model rotating SinSynaptosome.pdb and SinSynaptosome_symmetry.pdb – updated model for the Sin synaptosome. Each file contains one *res* site and the proteins bound to it. The full synaptosome consists of both files. Note that each file also contains the symmetry information necessary to regenerate the other.

Tn3synaptosome.pdb and Tn3synaptosome_symmetry.pdb – new model for the Tn3 synaptosome Each file contains one *res* site and the proteins bound to it. The full synaptosome consists of both files. Note that each file also contains the symmetry information necessary to regenerate the other.

Table 1 – crystallographic statistics

---

**Figure 1 supplement 1.**
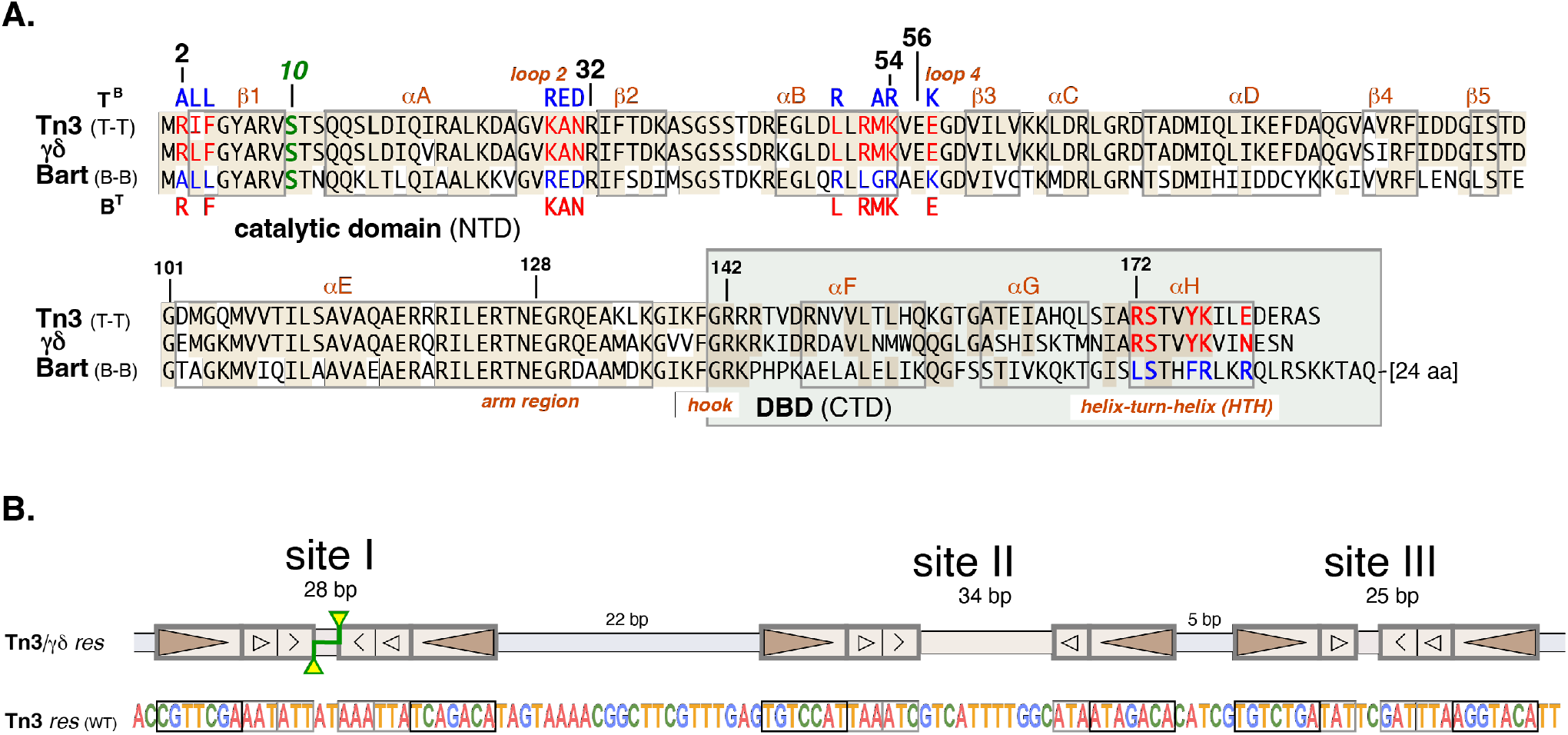
Relevant Sequences. a) Protein sequences. The Tn3 and γδ resolvases are 80% identical; Tn3 and Bart are only 49% identical (shaded residues). Bart has distinct *R* interface and DNA binding (DBD) specificities, and differs at relevant residues (highlighted, red/blue). The catalytic serine (S10), canonical *R* interface residues (2, 32, 54, 56), and E128 in helix E, are also highlighted. The chimaeric catalytic domains designated T^B^ (Tn3 domain, Bart-type *R* interface) and B^T^ (Bart domain, Tn3-type *R* interface) have ‘patch’ mutations at the 10 positions shown, as described (Rowland et al, 2020). The 24-residue CTD tail of the natural Bart resolvase was deleted for all experiments described here. b) Sequence and organization of the wild-type Tn3 *res* site. The filled triangles denote regions bound by the helix-turn-helix DBD, smaller open triangles denote regions bound by the AT hook, and arrow heads denote regions bound by the C-terminal portion of helix E. Yellow triangles at site I indicate DNA cleavage sites. c)

**Figure 2, figure supplement 1.**
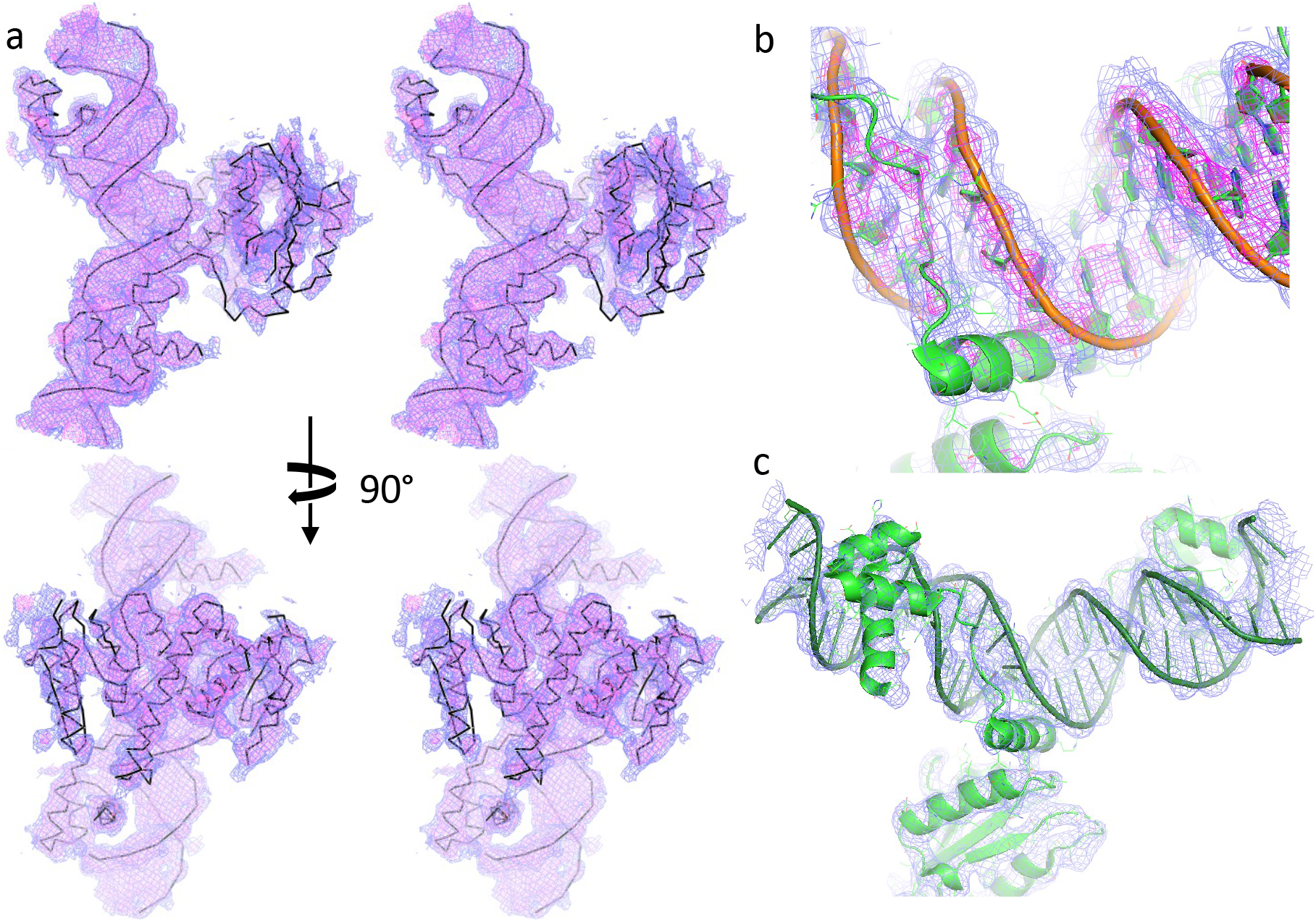
Electron density. a) Experimentally phased electron density map for crystal Form I. Two orthogonal stereo (wall-eyed) views were displayed in PyMOL and contoured at 1 sigma (blue) and 1.5 sigma (pink), with a “carve” radius of 4 Å to remove density for symmetry-related complexes. b) Final weighted 2Fo-Fc map for Form I, contoured at 1 and 3 sigma, with carve=4 Å. c) Final weighted 2Fo-Fc map for Form II, countered at 1.5 sigma, with carve = 4 Å.

**Figure 5 – supplement 1.**
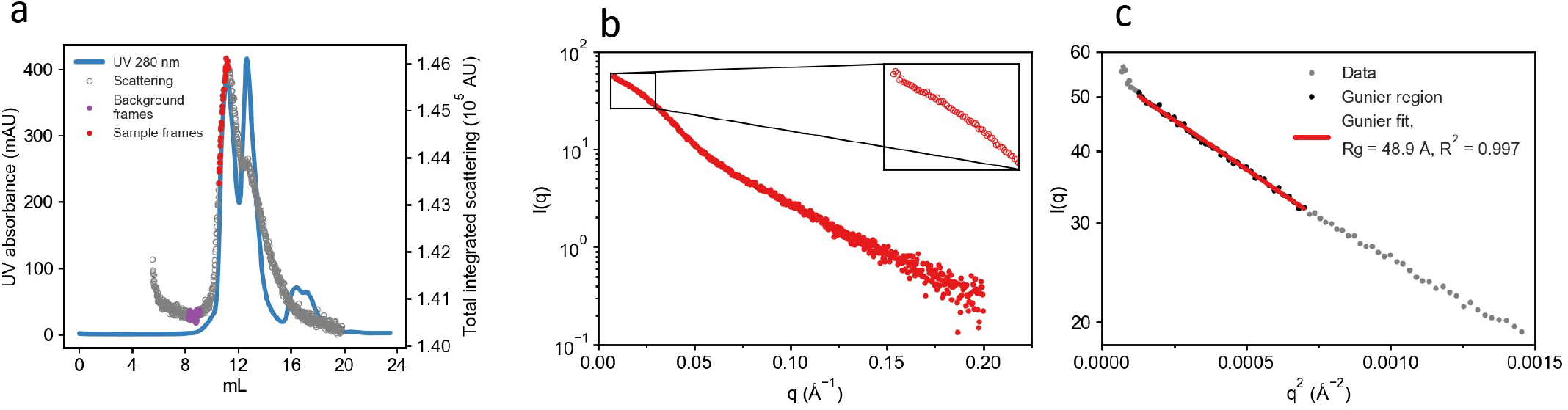
SEC-SAXS data for the Tn3 resolvase - site II complex. SAXS data were collected in-line following FPLC-driven size exclusion chromatography of a mixture of the Tn3 resolvase protein and a site II DNA fragment. a) Post-column FPLC UV A280 detector data (blue line) overlaid with the total integrated scattering from each SAXS exposure (circles). The 50 exposures marked as purple circles were merged for use as the background/buffer scattering reference and subtracted from the 40 exposures in red circles from the leading edge of the earliest-eluting peak. These buffer-subtracted data were then scaled and averaged to yield the final SAXS curve. b) The scaled/merged SAXS curve for the Tn3 resolvase - site II complex. The inset shows a magnification of the low q region. Data were truncated to q < 0.2 Å^-1^. c) Guinier fit and radius of gyration (Rg) determination. The SAXS data in the region between the first data point and q = 0.039 are displayed as points, transformed as a Guinier plot (log intensity as a function of q^2^). The Guinier region identified by AutoRG is given as black points, with the linear Guinier fit as a red line. The calculated Rg is 48.9 Å. In all cases, the scattering vector q is given as 4π sin θ/λ (Å^-1^).

**Figure 6 Supplement 1.**
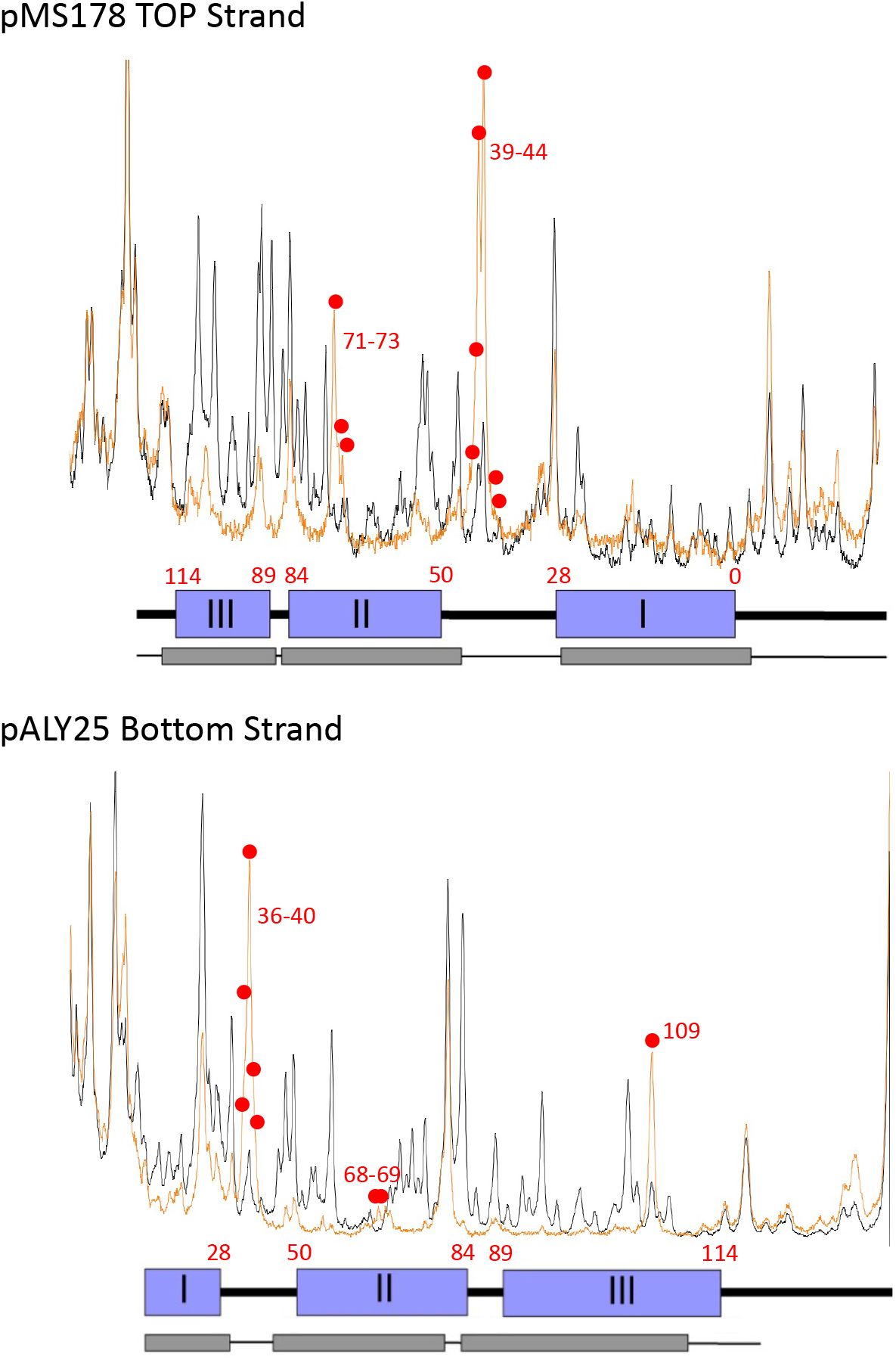
DNase I footprinting of the Tn3 synaptosome. The plots show representative data sets of DNase I strand cleavage intensity within *res*, for naked DNA (black lines) and DNA within the crosslinked synaptosome (orange lines). The positions on the x-axis corresponding to *res* binding sites are indicated by the blue boxes below the plots, and the thinner grey bars indicate regions of general protection from DNase I cleavage by resolvase in the synaptosome. Note that protection of the site I DNA is less complete than for sites II and III; we think that this is due to partial dissociation of site I-bound resolvase subunits during crosslinking of the synaptosomes prior to DNase I treatment ((Watson, 1994); see Materials and Methods section). The small red-filled circles indicate major enhancements of cleavage of the synaptosome DNA by DNase I at specific phosphodiester bonds.

**Figure 7 Supplement 1.**
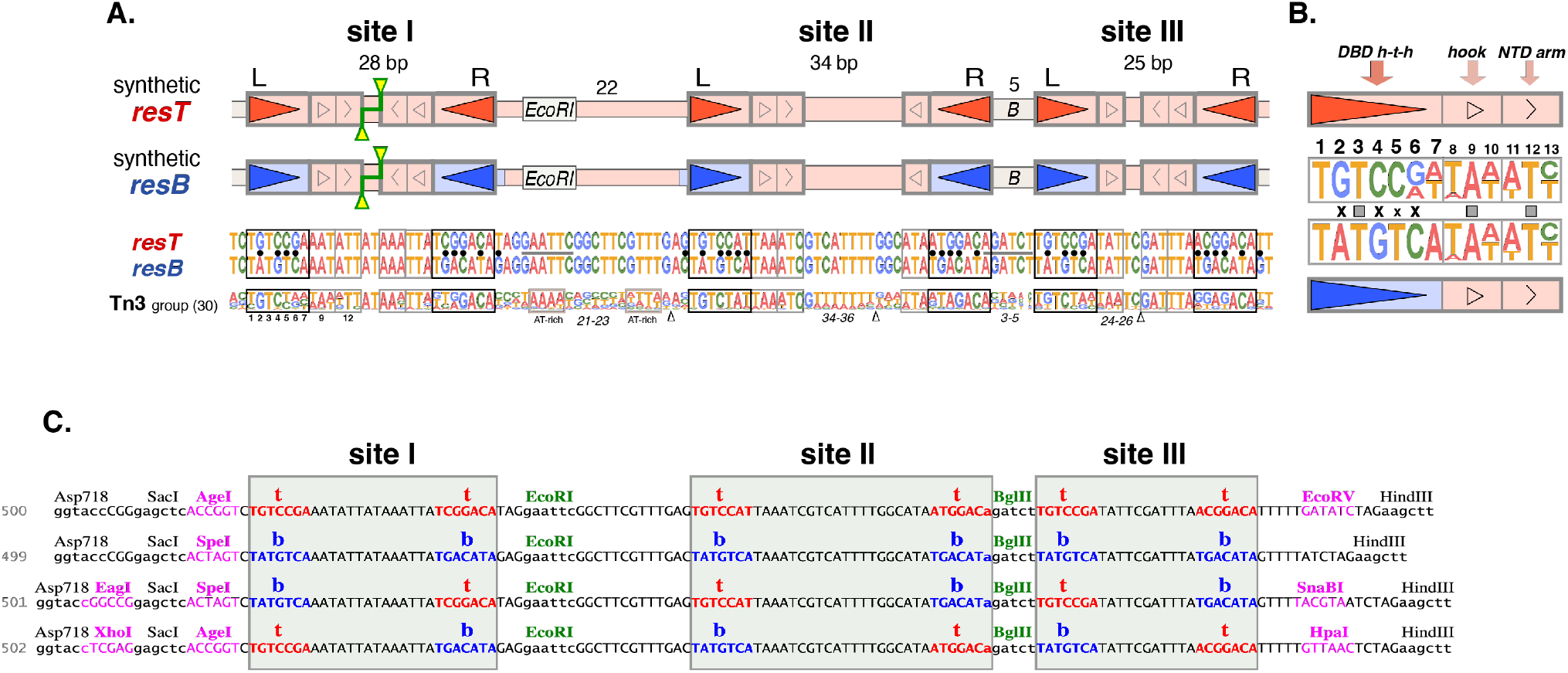
Design and sequence of synthetic *res* sites a) Design of synthetic *res* sites used for targeting experiments. The novel chimaeric *res* sites used here are all hybrids of two synthetic sequences, *resT* and *resB* (cognate sites for Tn3 and Bart resolvases), previously described (Rowland et al., 2020). Both sites are based on wild-type Tn3 *res* sequences (all segments shown in pale red). They differ in the 7 bp motifs (bold red/blue arrows) recognized by the helix-turn-helix part of the Tn3 or Bart DBD. Differences between *resT* and *resB* are highlighted (•) in the sequences. *resT* and *resB* retain many features of natural *res* sites – as exemplified by the *res* site logo based on 30 diverse Tn3-type sites - notably the highly non-random ‘linker’ sequences between the DBD motifs. (Rowland et al., 2020) b) CTD targeting motifs. Sequence logos compare the base frequencies (linear scale) in the six half-sites of *resT* and *resB* (excluding positions 11–13 of site IIR and site IIIL). Positions 2, 4, 5 and 6 are thought to be key determinants for selective DBD binding at the *resT* and *resB* motifs; the natural motifs were altered at ten positions to minimize potential selectivity overlaps (see Rowland *et al*, 2020). Note that the motifs contacted by the catalytic domain arm (the C-terminus of helix E) (positions 11-13, open arrows) and the DBD ‘hook’ (positions 8-10, triangles) are not thought to differ for the Tn3 and Bart resolvases. c) Synthetic recombination sites used in substrate constructions. Sequences 500 and 499 correspond to *resT* and *resB*, as previously described (Rowland et al., 2020). Sequences 501 and 502 correspond to the hybrid *res* sites (bt tb tb) and (tb bt bt). All other hybrid *res* sites were derived from these four synthetic sequences. The 7 bp motifs at the left and right ends of sites I, II and III are highlighted *red* (Tn3) and *blue* (Bart); within sites I, II and III, there are no further differences between these four sequences. Sites I and II are separated by an *Eco*RI restriction site, and sites II and III by a *Bgl*II site. Restriction sites in *magenta* were used as markers during plasmid construction. Substrate sequences are available on request.

**Figure 8 Supplement 1.**
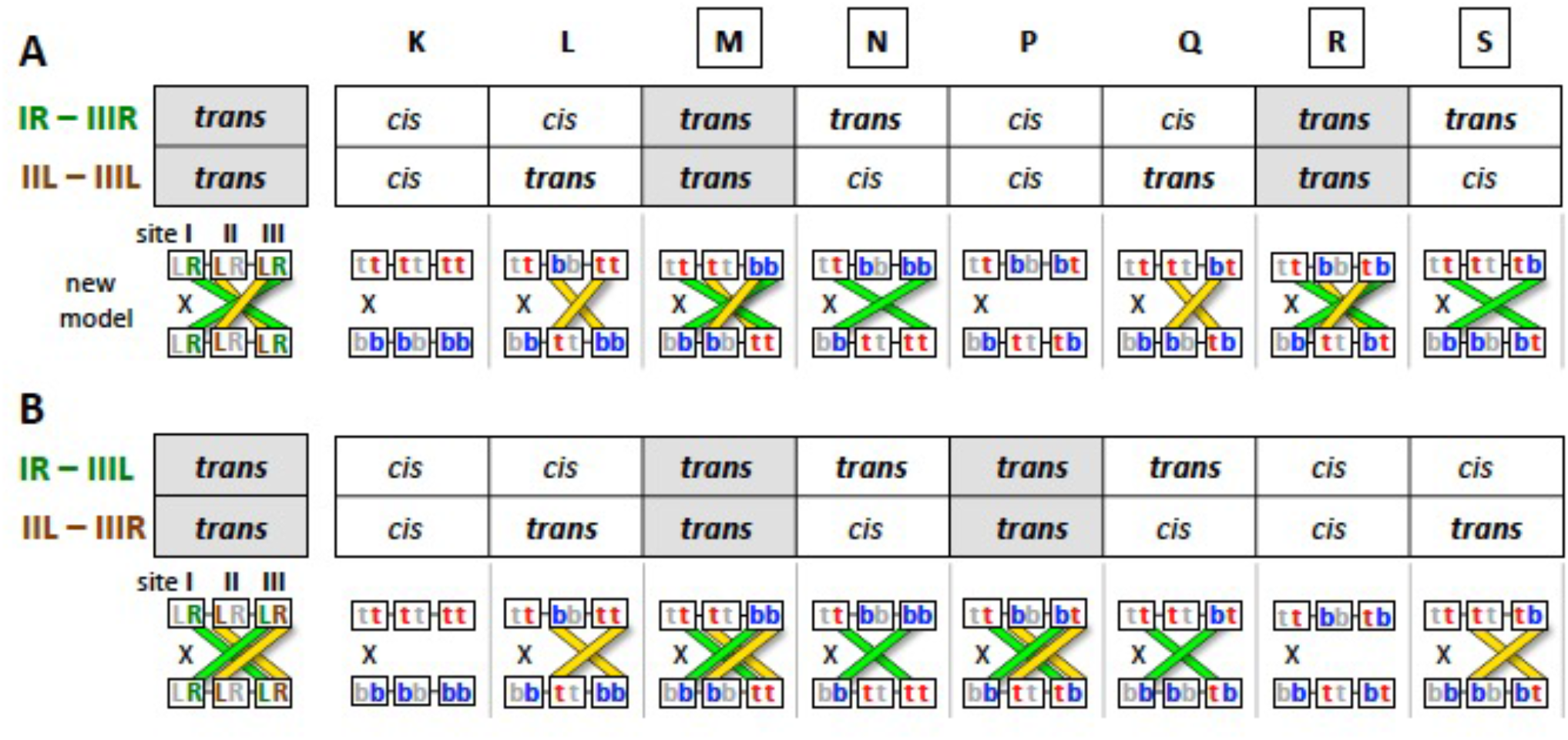
Further explanation of how the experiments presented in Figure 8, and the eight recombination substrates used, provide strong support for the new synapse model (Figure 6), and rule out alternative arrangements of the dimer-dimer *R* interfaces. Specifically, as shown here, the data rule out an alternative model in which the orientation and connectivity of the site III dimer are reversed (b). Similar arguments (not shown) help to rule out the six other possible permutations of *cis* or *trans* connections between subunits IR + IIIL and IIL + IIIR, or IR + IIIR and IIL + IIIL, including the two arrangements suggested by (Sarkis et al., 2001), with subunits IIL + IIIL paired *in cis*. a) This part is identical to Figure 8c and d, and shows, for each substrate, whether IR-IIIR and IILIIIL interactions between matching subunits can occur *in cis* or *in trans*. The cartoons show, for each substrate, only the half-sites that match *in trans*: IR-IIIR (*green*) and IIL-IIIL (*yellow*). The four substrates that are efficiently recombined, as shown in Figure 8b, are M, N, R and S (boxed). b) An alternative scenario in which the site III dimer is in the opposite orientation and the interactions are IR-IIIL and IIL-IIIR (both *in trans*). Here, the predicted requirements in order for half-sites to match *in trans* do not correlate with the experimental data.

**Figure 10 supplement 1.**
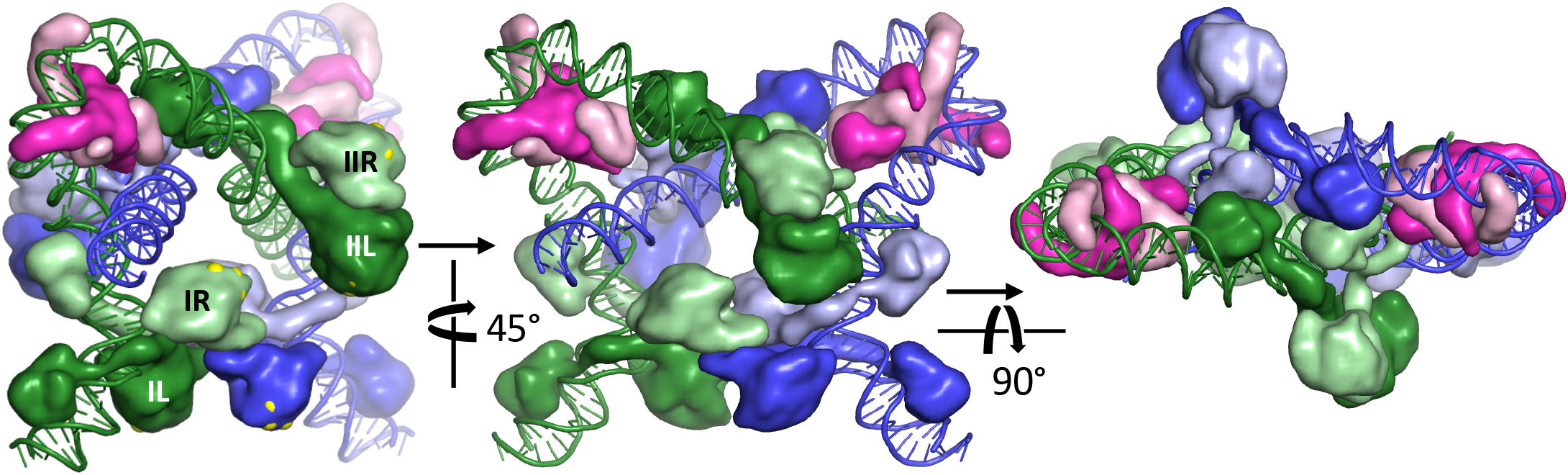
Updated model for the Sin Synaptosome. Three views of the model are shown, in orientations similar to those shown for the Tn3 synaptosome in the main Figure 6. This model differs from that previously published (Mouw et al., 2008) primarily in that the site I-bound subunits are based on a constitutively active Sin rather than γδ resolvase tetramer (see Materials and Methods). Proteins are shown as smoothed surfaces, IHF heterodimers are in pink, and Sin subunits are colored according to the DNA segment they are bound to, with those bound to the right half of each site in a lighter shade than those bound to the left. In the left panel, each protein is labeled according to which half site it is bound to. Also in the left panel, the Cα atoms of *R* interface residues (F52, R54 and D57) are shown as large yellow balls poking out of the protein surface. This model, based on rigid-body docking of substructures, does not perfectly recapitulate the *R* interface between the site I-right and site II-left bound proteins. However, relatively modest adjustments, mostly at the kinked points in the site II-bound proteins’ E-helices, would make it do so. Note that in the Sin case, the central node is held together by contacts between the DBDs of the site II-bound proteins (most obvious in the right panel).

