## Supplementary material for "Structural basis for topological regulation of Tn3 resolvase": Table 1. Crystallographic Data Collection and Refinement Stastics.

Table 1. Data collection and refinement statistics.

|  | Form I, PDBid 5cy1  SBgrid data set #682  1 dimer / asymmetric unit | Form II, PDBid 5cy2  SBgrid data set #683  2 dimers / asymmetric unit |
| --- | --- | --- |
| Wavelength | 0.97948 | 0.97948 |
| Resolution range | 29.19 - 3.4 | 49.08 - 4.0 (4.143 - 4.0) |
| Space group | P 43 21 2 | C 1 2 1 |
| Unit cell | 77.31 77.31 272.28 90 90 90 | 144.9 151.92 106.14 90 99.85 90 |
| Total reflections | 92434 | 67699 |
| Unique reflections | 12096 (1185) | 19086 (1812) |
| Multiplicity | 7.6 (6.7) | 3.5 (2.6) |
| Completeness (%) | 1.00 (0.998) | 0.99 (0.95) |
| Mean I/sigma(I) | 34.2 (1.5) | 22.9 (1.0) |
| Wilson B-factor | 146.49 | 182.64 |
| R-merge | 5.7% (> 100%) | 8.7% (>100%) |
| CC1/2 | (0.636) | (0.486) |
| Reflections used in refinement | 12091 (1184) | 19058 (1794) |
| Reflections used for R-free | 575 (60) | 968 (100) |
| R-work | 0.2509 (0.3782) | 0.2072 (0.3376) |
| R-free | 0.3002 (0.4409) | 0.2497 (0.3646) |
| Number of non-hydrogen atoms | 4072 | 7792 |
| macromolecules | 4072 | 7792 |
| Protein residues | 363 | 713 |
| RMS(bonds) | 0.003 | 0.010 |
| RMS(angles) | 0.54 | 1.09 |
| Ramachandran favored (%) | 96 | 96 |
| Ramachandran allowed (%) | 4.2 | 3.9 |
| Ramachandran outliers (%) | 0 | 0.43 |
| Rotamer outliers (%) | 0.33 | 0.51 |
| Clashscore | 4.53 | 6.13 |
| Average B-factor | 188.70 | 221.00 |
| macromolecules | 188.70 | 221.00 |
| Number of TLS groups | 9 | 16 |

Statistics for the highest-resolution shell are shown in parentheses.
